# Keratin degradation reflects a starvation survival strategy in *Fervidobacterium islandicum* AW-1

**DOI:** 10.1101/2025.07.04.663136

**Authors:** Jae-Yoon Sung, Ji-Yeon Kim, Hyeon-Su Jin, Je-Hyun Baek, Nicole Enjeh Kim, Hyun Ho Song, Seong-Hun Bong, Yong-Jik Lee, Byoung-Chan Kim, Do Yup Lee, Dong-Woo Lee

**Author notes:** Corresponding authors, **Do Yup Lee**, Department of Agricultural Biotechnology, Center for Food and Bioconvergence, Research Institute for Agricultural and Life Sciences, Seoul National University, Seoul 08826, South Korea., **Dong-Woo Lee**, Department of Biotechnology, Yonsei University, Seoul 03722, South Korea. These authors contributed equally to this work (Jae-Yoon Sung, Ji-Yeon Kim).

## Abstract

Keratin is a highly cross-linked, disulfide-rich protein that resists proteolysis, which poses a major challenge for microbial degradation. Here, we show that *Fervidobacterium islandicum* AW-1 initiates a starvation-induced keratinolytic program involving membrane-associated proteases and redox-mediated sulfitolysis. Multi-omics integration reveals that nutrient limitation triggers global metabolic reprogramming, promoting sulfur assimilation, biofilm formation, and chemotaxis-linked persister-like adaptation. Substrate-specific transcriptomics identified a temporally regulated protease repertoire tightly coordinated with sulfitolytic activity, facilitating efficient feather decomposition under starvation. Protein-protein interaction networks uncovered stress-responsive transcriptional regulators that govern this process. Time-resolved gene expression analysis and metabolomic profiling further revealed that cyclic-di-GMP signaling, stringent response, and flagella assembly mediate transitions between motility and sessile growth, contributing to surface colonization and persistence. Together, our findings establish a starvation-responsive survival mechanism that couples keratin degradation to stress adaptation in extreme environments, offering insights into microbial persistence and potential strategies for keratin valorization.

---

Microbial survival under nutrient scarcity relies on a range of adaptive strategies, including sporulation, biofilm formation, and metabolic plasticity ^1, 2^. These adaptations have enabled prokaryotes to persist in extreme environments for over 3.2 billion years, shaping their ecological success and evolutionary resilience ^3, 4^. For example, Gram positive Bacillota (formerly Firmicutes) undergo sporulation to enter dormant states under starvation ^5^, while Gram negative Pseudomonadota (Proteobacteria) activate chemotaxis and biofilm formation to enhance nutrient acquisition and stress tolerance ^6^. Biofilm formation confers structural protection and increases resistance to environmental stressors and antibiotics, especially in pathogenic bacteria ^7^. Starvation also triggers the formation of persister cells, generating a slow-growing subpopulation that exhibits extreme tolerance to stress, including antibiotics ^8^. Unlike stationary-phase cells, persisters are metabolically active at minimal levels, allowing survival under prolonged nutrient deprivation ^9^.

Metabolic plasticity further supports microbial survival by enabling the utilization of recalcitrant biopolymers such as lignocellulose, chitin, and keratin as alternative nutrient sources ^10^. Among these, keratin is one of the most degradation-resistant due to its dense hydrogen bonding and extensive disulfide cross-linking ^11^, forming a fibrous matrix that provides structural integrity to tissues such as skin, feathers, hair, and hooves. While its resistance poses a barrier to degradation, certain microorganisms, particularly bacteria and fungi, have evolved specialized enzymatic systems to degrade keratin, allowing survival in oligotrophic environments ^12, 13, 14^.

Previously, we identified *Fervidobacterium islandicum* AW-1, an extremely thermophilic anaerobe of the order Thermotogales, capable of degrading native keratin at 70°C ^15, 16, 17^. Unlike canonical extracellular proteolysis, keratin degradation by *F. islandicum* AW-1 involves a dual mechanism of membrane-associated proteolysis and redox-mediated sulfitolysis ^18^. Sulfitolysis, which cleaves disulfide bonds in keratin, facilitates access to peptide backbones and accelerates degradation. Prior studies revealed the upregulation of membrane-associated metalloproteases and cytosolic peptidases under starvation, suggesting that keratinolysis functions as a stress-adaptive process rather than a constitutive nutrient response ^12, 19^. Similar adaptations are found in dermatophytic pathogens, where secreted and membrane-bound peptidases mediate keratin breakdown during host colonization ^20^. Furthermore, keratin degradation has been linked to sulfur metabolism, including Fe-S cluster biogenesis and thiol generation, highlighting the role of redox regulation in enabling proteolysis under stress ^18, 21^.

Despite these insights, fundamental questions remain: What molecular signals trigger keratinolytic activation? How do metabolic networks reorganize to support keratin utilization during starvation? And what regulatory circuits govern this complex response? To address these questions, we employed a systems-level, genome-wide multi-omics approach combining transcriptomics, proteomics, and metabolomics to dissect starvation-induced keratinolysis in *F. islandicum* AW-1.

Our integrated analysis reveals that keratin degradation is a starvation-induced survival mechanism, rather than a substrate-specific response. Nutrient limitation triggers membrane-associated keratinolysis, stress-responsive gene expression, and metabolic rewiring, including activation of sulfur assimilation and amino acid catabolism. We also identify a dynamic transition from sessile biofilm formation to motility-driven dispersal through swarming—a collective behavior mediated by flagella synthesis and cyclic-di-GMP signaling. This phenotypic plasticity enables *F. islandicum* AW-1 to efficiently colonize and degrade insoluble keratin substrates, ensuring persistence in keratin-rich, soluble nutrient-depleted environments.

By integrating time-resolved multi-omics data, this study provides a mechanistic model of how keratin degradation functions as a broader metabolic adaptation to environmental stress. These findings offer new insight into microbial survival in extreme environments and suggest potential biotechnological applications for keratin waste valorization and biofilm-related stress tolerance.

## Keratin degradation by *F. islandicum* AW-1 is redox-enhanced but not substrate-induced

To determine whether *F. islandicum* AW-1 can utilize keratin as a nutrient source, we monitored growth and amino acid release during anaerobic cultivation with native feathers at 70°C. Feather-grown cells exhibited sustained proliferation, progressive degradation of feather biomass, and accumulation of amino acids consistent with keratin degradation, including Ala, Gly, Ser, Val, and Ile, as well as essential amino acids (Met, Trp, and Lys) (**Table S1 and Fig. S1a–b**). Similar levels of fermentation end-products, including lactate (6 mM) and acetate (3 mM), were observed in both glucose- and feather-supplemented cultures (**Fig. S1c**), confirming that keratin serves as a viable carbon and nitrogen source.

To evaluate nutrient-dependent effects on keratinolysis, we compared growth and protease activity across various carbon/nitrogen sources. Cultures grown in glucose, peptone, tryptone, or casein supported rapid early-phase growth, while feather-grown cells showed slower onset but reached comparable or higher final cell densities, accompanied by increased accumulation of free amino acids (**Figs. 1a and S1d)**. Total protease activity, measured using casein as a substrate, was comparable across all conditions (**Fig. 1b**), and feather hydrolysis by crude cell extracts showed only minor differences under reducing conditions (**Fig. 1c)**. Varying feather concentrations also had minimal impact on keratinolytic activity (**Fig. S1e–g**), suggesting that keratin does not transcriptionally induce a dedicated or specialized protease response.

**Figure 1.**
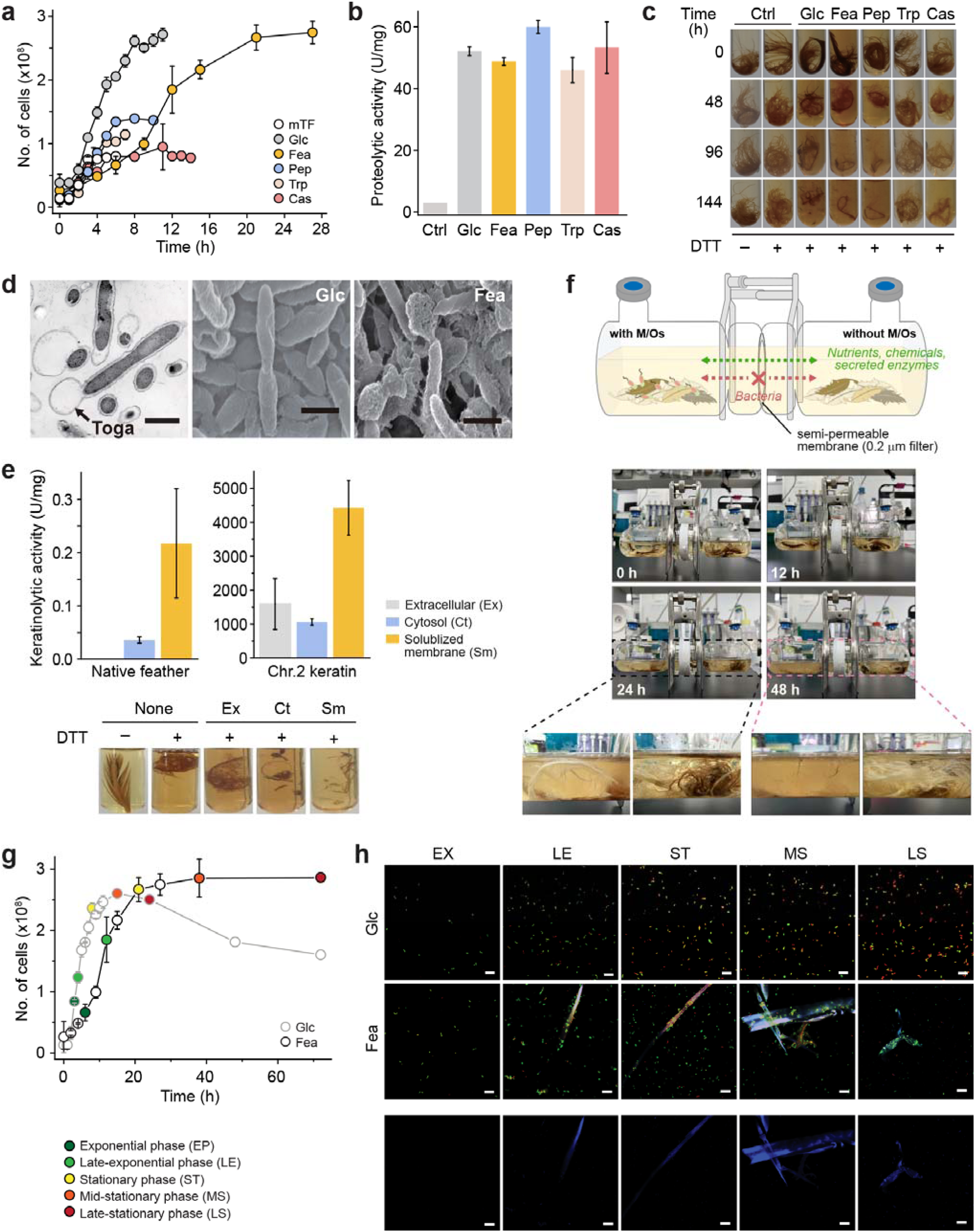
Growth physiology and keratin degradation by *F. islandicum* AW-1. (**a**) Growth curves in modified TF (mTF) medium with glucose (Glc), peptone (Pep), tryptone (Trp), casein (Cas), or native feathers (Fea) at 70 . (**b**) Total proteolytic activity in crude extracts (0.1 mg/ml) from late-exponential phase cells. (**c**) Feather degradation by crude extracts with and without 10 mM DTT at 75°C. **(d)** SEM images showing Glc-grown rods and Fea-grown cells with balloon-like toga morphology (scale bar = 1 µm). **(e)** Keratinolytic activity of subcellular fractions from Fea-grown cells for 24 h incubation using native feathers (left) or recombinant Chr. 2 keratin ^48^ (right) as substrates. **(f)** Two-chamber assay confirming cell-associated hydrolysis. **(g)** Growth curves in Glc-vs. Fea-supplemented media. **(h)** CLSM images of cells during growth on glucose or feathers stained with SYTO9 (green, live) and PI (red, dead); autofluorescent feather fragments stained with CPM. Scale bars = 10 µm.

To test whether redox conditions influence keratin degradation, we examined amino acid release in the presence of dithiothreitol (DTT), a reducing agent that promotes disulfide bond cleavage. DTT significantly enhanced feather solubilization and amino acid liberation (**Figs. 1c and S1d, f–g**), indicating that sulfitolysis—thiol-mediated disruption of disulfide crosslinks—facilitates proteolytic access to keratin’s rigid backbone, potentially associated with redox metabolic changes.

Together, these findings suggest that *F. islandicum* AW-1 constitutively expresses keratinolytic enzymes, but keratin itself does not induce their expression. Instead, degradation efficiency is enhanced under reducing conditions via redox-mediated sulfitolysis, enabling efficient keratin utilization during nutrient limitation.

## Membrane-associated keratinases drive proteolysis under nutrient scarcity

To investigate cellular adaptations under nutrient limitation, we examined cell morphology using scanning electron microscopy (SEM). Glucose-grown *F. islandiucm* AW-1 cells maintained a smooth rod-shaped appearance, whereas feather-grown cells exhibited expanded outer envelopes characteristic of the Thermotogales “toga” structure (**Fig. 1d**). This starvation-induced membrane remodeling likely enhances cell-surface interaction with insoluble and hydrophobic keratin substrates. Consistent with this, fatty acid profiling by gas chromatography-flame ionization detection (GC-FAME) showed that feather-grown cells exhibited a shift from saturated palmitic acid (C_16:0_; 79% in glucose-grown cells) to increased unsaturated oleic acid (C_18:1_ *cis*-9, 23%) (**Table S2**), suggesting elevated membrane fluidity and flexibility under oligotrophic conditions.

To identify the subcellular localization of keratinolytic activity, we fractionated cells into extracellular, cytosolic, and solubilized membrane (SM) components. While extracellular protease activity was negligible, robust keratinolytic activity was enriched in membrane-associated fractions, particularly in feather-grown cells (**Figs. 1e and S1h**). These fractions efficiently degraded both native feathers and recombinant keratin substrates, establishing the SM proteome as the principal site of proteolysis under nutrient stress.

To assess whether keratin degradation is mediated by secreted or cell-associated enzymes, we employed a custom two-chamber system with a semi-permeable membrane barrier (**Fig. 1f**). Keratin degradation occurred only in chambers containing *F. islandicum* AW-1 cells, indicating that the initiation of keratin hydrolysis requires direct physical contact and is mediated by cell-bound enzymes rather than diffusible proteases or reducing agents such as cysteine and sulfite.

Confocal laser scanning microscopy (CLSM) further revealed shifts in spatial behavior and viability under nutrient limitation. Glucose-grown cells remained planktonic and lost viability during the stationary phase, whereas feather-grown cells formed dense sessile aggregates on feather surfaces, maintaining viability over prolonged incubation (**Figs. 1g and 1h**). These biofilm-like structures may facilitate localized proteolysis and resource acquisition.

Collectively, these findings indicate that *F. islandicum* AW-1 adopts a starvation-responsive strategy centered on membrane remodeling, protease localization, and surface colonization to enable efficient degradation of recalcitrant keratin under oligotrophic conditions.

## Nutrient-dependent transcriptomic profiling reveals sulfur metabolism, energy optimization, stress adaptation, and membrane-associated proteolysis in feather-grown cells

To uncover transcriptional adaptations underpinning keratin utilization, we performed RNA-Seq analysis of *F. islandicum* AW-1 grown under exponential-phase conditions (6-8 h) on native feathers, glucose, peptone, or tryptone (**Table S3**). Of 1,949 annotated protein-coding genes (GenBank accession NZ_CP014334.1) ^12^, 336 genes were differentially expressed in feather-grown cells relative to glucose, including 291 upregulated and 45 downregulated transcripts (fold change ≥2, *p* < 0.05) (**Figs. 2a, 2b, S2a, and DataSet S1)**.

**Figure 2.**
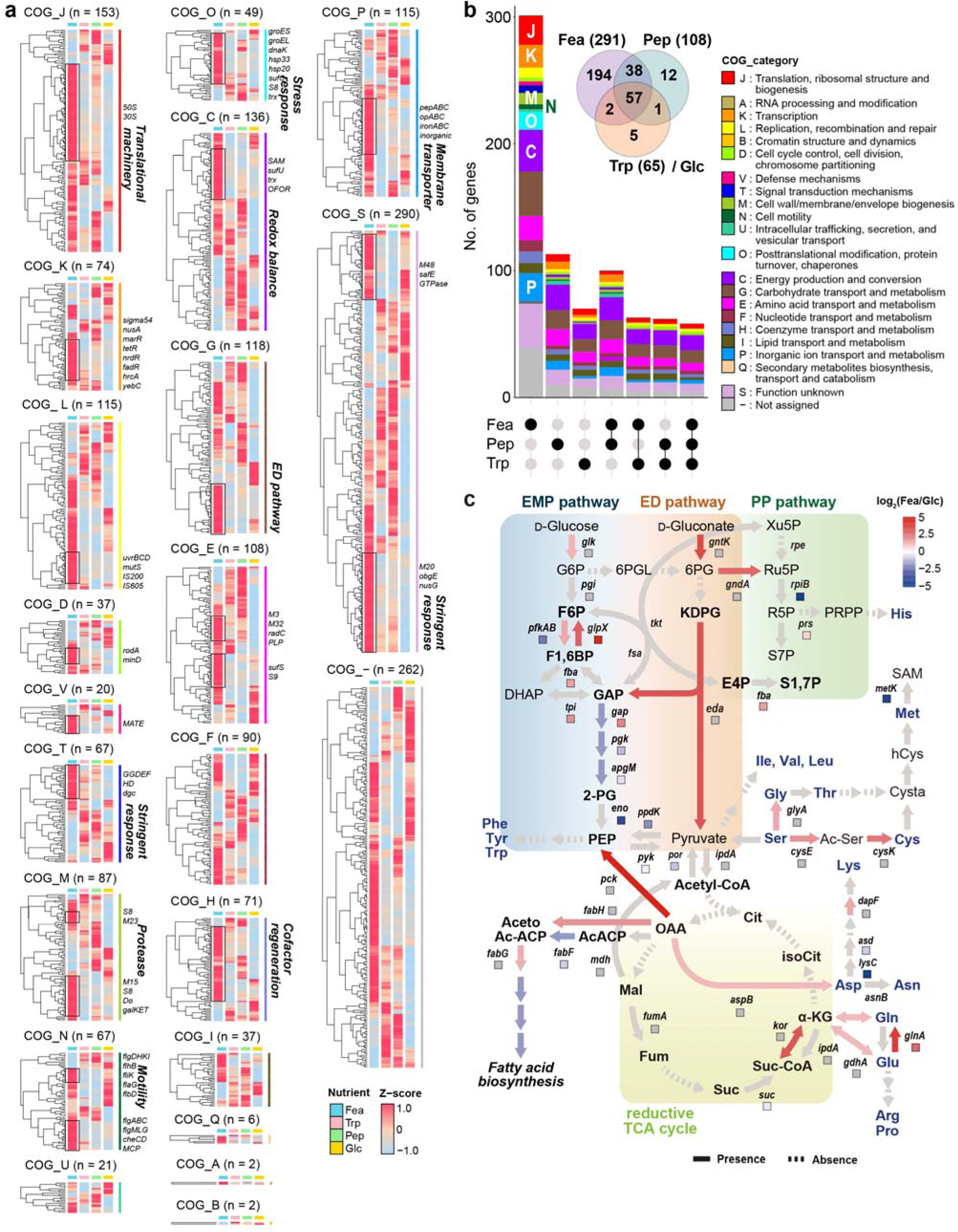
Nutrient- and time-dependent transcriptomic reprogramming in *F. islandicum* AW-1. (**a**) Heatmaps of differentially expressed genes (DEGs) (Fea, Pep, Trp vs. Glc at 8 h). DEGs are grouped by COG functional categories and hierarchically clustered based on Z-score normalized transcript abundance. Selected DEGs involved in sulfur metabolism (e.g., *suf*, *trx*, *isc*), chemotaxis, redox regulation, and stress response are annotated. **(b)** Functional classification of DEGs by COG. Bar plots show the number of up- and downregulated DEGs under each nutrient condition. The Upset plot (bottom) depicts shared and condition-specific DEGs, while the Venn diagram (top right) highlights unique and shared DEGs. **(c)** Metabolic pathway mapping showing altered flux in the Embden-Meyerhof-Parnas (EMP; blue), Entner-Doudoroff (ED; red), pentose phosphate (PP; green), and reductive TCA cycles (yellow). Amino acid biosynthesis pathways (e.g., Ser, Cys, Glu, Pro, Asn) are highlighted. The dotted arrows indicate reactions absent in the *F. islandicum* AW-1 genome. Arrows and squares represent transcriptomic and proteomic data, respectively. Colors indicate log FC of expression under Fea compared to Glc conditions (*p* < 0.05).

Functional categorization revealed transcriptional enrichment in sulfur metabolism, redox regulation, membrane-associated proteolysis, nutrient transport, and stress response pathways (**Figs. 2a, 2b, and S2b**). Central carbon flux was rewired toward the Entner–Doudoroff (ED) pathway, as evidenced by the upregulation of 6-phosphogluconate dehydratase and KDPG aldolase, alongside the downregulation of glycolytic phosphofructokinase (PFK). This transcriptional shift may favor ATP-sparing metabolism suited to oligotrophic environments (**Figs. 2c and S2b and DataSet S1**). Genes encoding reverse tricarboxylic acid (TCA) cycle enzymes, including ATP citrate lyase and two ferredoxin-dependent oxidoreductases, were also upregulated, indicating biosynthetic flux redistribution under low-energy conditions. The induction of the tripartite tricarboxylate transporters (TTTs) suggests scavenging of organic acids, likely derived from keratin hydrolysates (**Fig. 2c**).

A major transcriptional feature was the pronounced activation of sulfur assimilation and redox-balancing systems. These included the sulfur formation (Suf) system (e.g., *sufB*, *sufC*), cysteine synthase, radical SAM enzymes, and Fe-S cluster assembly proteins (**Figs. 2 and S2a–b**). Together, these findings support a model in which upregulated sulfur metabolism may confer to redox-mediated sulfitolysis, potentially facilitating disruption of keratin’s disulfide structure and maintaining redox homeostasis.

Stress-related transcriptional regulators were also strongly modulated. TetR-, MarR-, and FadR-family repressors were induced >24-fold, while the NADH-sensing repressor Rex was repressed (**DataSet S1: Sheet 1**). Concurrent induction of molecular chaperones (e.g., DnaK, GroES), DNA repair proteins, and heat shock proteins indicates proteotoxic and oxidative stress during keratin catabolism (**Figs. 2a, 2b and S2a**). Nutrition acquisition systems, including ABC transporters for amino acids and peptides, solute-binding proteins, and inorganic ion channels, were also upregulated. In contrast, ATP-intensive processes such as thiamin biosynthesis (*thiC*, *thiS*, *thiE*), and heavy metal efflux were downregulated (**Figs. 2 and S2a–b; DataSet S1: Sheet 1**), reflecting energy prioritization toward survival and catabolism.

Consistent with membrane restructuring, genes encoding membrane-bound metalloproteases (e.g., M23 and M48), cytosolic peptidases (e.g., M38, M55, and M16), and carboxypeptidases (e.g., C15 and M32) were significantly induced, although with moderate absolute expression (**Fig. S2a and S2c; DataSet S1**). These enzymes likely contribute to keratin turnover, proteostasis, and redox-modulated proteolysis. Upregulation of chemotaxis and motility-related genes (e.g., *fliC*, *flhA*, methyl-accepting chemotaxis proteins) further supports a coordinated behavioral response for surface colonization, motility, and substrate engagement (**Figs. 1f, 1g, 2a and S2a**).

Importantly, 194 DEGs were uniquely enriched in feather-grown cells, excluding overlap with peptone- or tryptone-grown conditions (**Fig. 2b and DataSet S1: Sheets 1-3**). These feather-specific genes include keratinolytic proteases, sulfur metabolism enzymes, redox regulators, and metabolic nodes (e.g., ED and reverse TCA) tailored to low-energy, disulfide-rich environments.

In summary, *F. islandicum* AW-1 engages a multi-tiered transcriptional response during keratin utilization, characterized by (i) sulfur-driven sulfitolysis, (ii) energy-optimized metabolism, (iii) membrane-associated proteolysis, and (iv) stress-resilient transport and regulation. This network, orchestrated by global regulators including MarR, TetR, and Rex, exemplifies a microbial persistence strategy adapted to keratin-dominated, oligotrophic ecosystems.

## Time-resolved transcriptomic analysis reveals a transition to sessile adaptation and stress resilience during keratin degradation

To delineate temporal regulatory dynamics during keratin degradation, we performed RNA-Seq analysis of *F. islandicum* AW-1 grown on native feathers at 8 h and 12 h (**Fig. 3a–3b**). Among 57 predicted protease genes, ∼60% exhibited peak Z-score normalized expression in feather-grown cells, though most maintained modest transcript abundance (<1000 TPM). Notably, one S8-family protease maintained high expression (>4000 TPM), but overall, protease expression reflected a nutrient-responsive rather than keratin-specific induction pattern (**Fig. S2c**).

**Figure 3.**
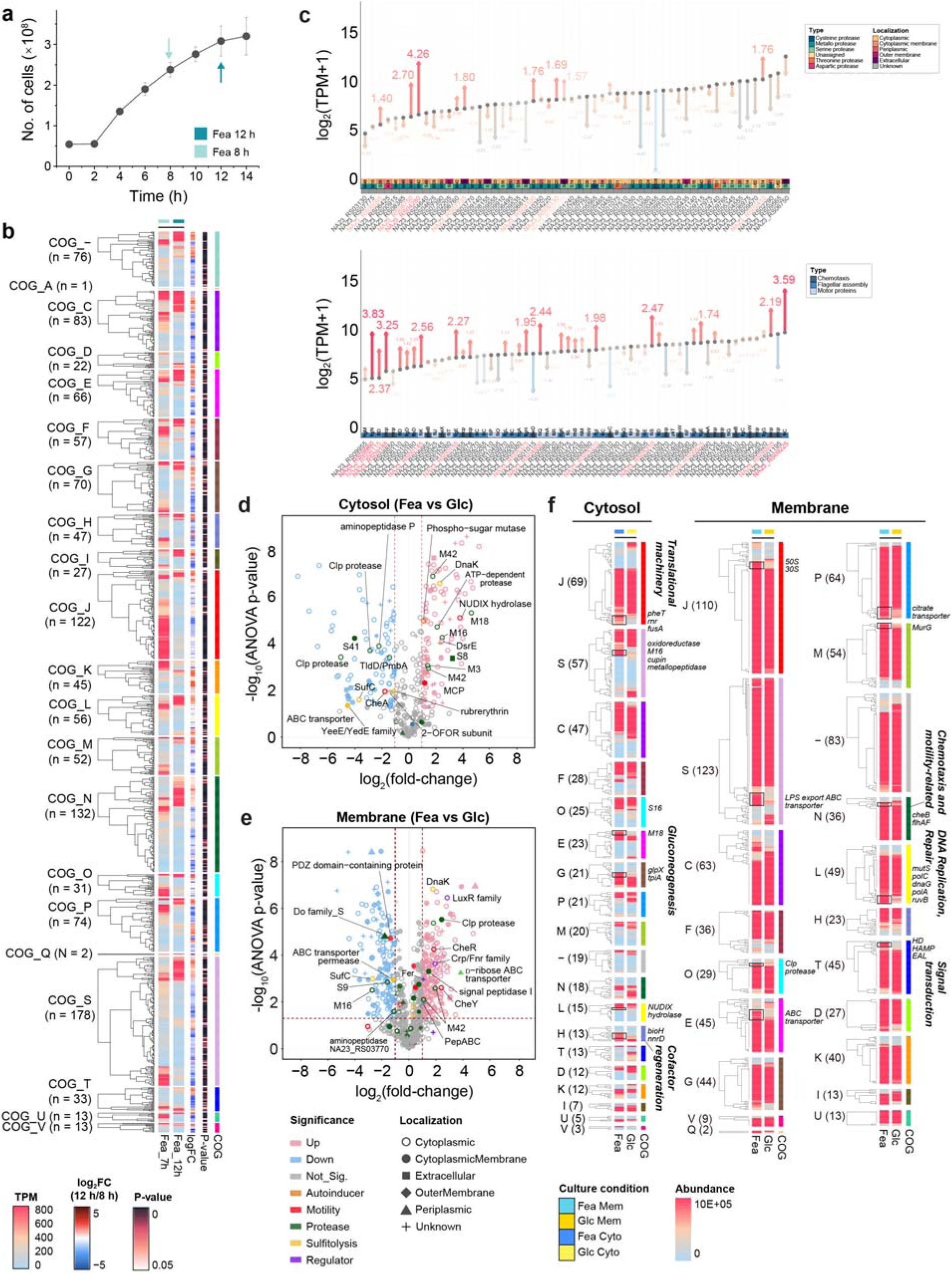
Temporal transcriptomic and subcellular proteomic profiling of keratinolytic responses in *F. islandicum* AW-1. **(a)** Growth curve of *F. islandicum* AW-1 in mTF medium supplemented with feathers. Arrows indicate RNA-Seq sampling points at 8 h and 12 h. **(b)** Heatmaps of time-dependent transcriptomic changes in feather-grown cells, categorized by COG functional classes. Left panel: Z-score normalized expression levels at 8 h; center panel: log₂ fold changes at 12 h; right panel: adjusted *p*-value. (**c**) Gene expression profiles of protease-related genes (upper panel) and chemotaxis/motility-related genes (lower panel) comparing 12 h to 8 h in feather-grown cells. Expression values are shown as log₂(TPM + 1), with red and blue arrows indicating upregulated and downregulated genes, respectively. (**d**, **e**) Volcano plots of differentially expressed proteins (DEPs) in cytosolic (**d**) and membrane (**e**) fractions from feather-versus glucose-grown cells. Significantly upregulated proteins (log_2_FC > 1, *p* < 0.05) are shown in red, downregulated (log_2_FC > 1, *p* < 0.05) in blue, and non-significant in gray. (**f**) Heatmaps of Z-score normarlized abundances for DEPs from cytosolic and membrane compartments, categorized by COG functional groups and growth conditions.

Between 8 h and 12 h—corresponding to a transition from active growth to stress adaptation—we observed distinct reprogramming of protease expression profiles (**Figs. 3c and S3**). Proteases associated with stress responses (e.g., M42, M48, and S14) were upregulated >8-fold, while intracellular protein quality control enzymes (e.g., M15 and M38) also showed >3-fold induction. In contrast, most other protease transcripts declined, indicating a functional reshaping of the proteolytic landscape. This proteolytic transition was accompanied by repression of chemotaxis regulators (*cheA*, *cheC*, and *cheD*), while flagellar genes, including *fliC* (flagellin), were upregulated, suggesting a behavioral shift from environmental sensing to surface-associated motility, likely facilitating substrate adhesion.

Redox and sulfur metabolism genes (e.g., glutaredoxin, cupin, rubredoxin, Fur, and TauE) were broadly downregulated from 8 h to 12 h. The result suggested reduced transcriptional regulation in sulfur and redox systems as keratin hydrolysis progressed (**Figs. 3c and S3; Dataset S1: Sheet 9**). An exception was *safE*, encoding a sulfite exporter, which was upregulated >4-fold, possibly to eliminate excess reductants or sulfitolytic byproducts.

Genes involved in the stringent response (*obgE* and *hflX*) were repressed >10-fold, indicating reduced translational activity and ribosomal maintenance. Concurrently, cyclic-di-GMP turnover enzymes (e.g., EAL/GGDEF and HD-GYP domain proteins) were upregulated 2–5 fold, consistent with biofilm initiation and sessile adaptation (**Fig. S3**). This transition was further supported by global downregulation (>10-fold) of ribosomal protein genes, indicating entry into a metabolically quiescent, persister-like state (**DataSet S1: Sheet 9-12**).

Collectively, these findings reveal a coordinated transcriptional shift between 8 h and 12 h, wherein *F. islandicum* AW-1 transitions from active keratin metabolism to a metabolically quiescent, stress-tolerant state. This phase is characterized by reduced sulfur and redox activity, suppressed chemotaxis and ribosome biogenesis, and increased expression of stress-responsive proteases, flagellar components, and cyclic-di-GMP signaling proteins. These regulatory features highlight a survival-oriented program adapted for persistence in keratin-rich, nutrient-depleted environments.

## Proteome-wide analysis reveals nutrient-responsive enrichment of membrane-associated keratinases and motility proteins

To characterize protein-level adaptations during keratin utilization, we performed quantitative LC-MS/MS analysis of cytosolic and membrane-associated fractions from *F. islandicum* AW-1 grown on glucose or native feathers, focusing on both cytosolic and membrane-associated protein fractions. This approach allowed us to assess nutrient-dependent expression changes across subcellular compartments, particularly the enrichment of keratinolytic enzymes and stress-responsive regulators under feather-based growth.

Proteomic profiling identified a total of 910 proteins in feather-grown cells and 430 in glucose-grown cells, with a pronounced increase in membrane-associated proteins under keratin-based growth (**Fig. 3d–3f; DataSet S2**). In feather-grown cells, membrane fractions showed strong enrichment of ABC-type substrate-binding proteins, signal transduction components (CheR, Crp/Fnr family), and multiple proteases, including M42 family metallopeptidases and serine protease (**Fig. 3e–3f**). These proteases were largely undetectable or minimally expressed in glucose-grown cells, suggesting their induction is nutrient-dependent and associated with surface-bound keratin degradation.

In parallel, cytosolic fractions from feather-grown cells exhibited increased abundance of enzymes involved in amino acid fermentation, glutamate metabolism, and the gluconeogenesis, reflecting metabolic rewiring to support survival in oligotrophic conditions (**Fig. 3d and 3f**). Proteostasis-related chaperones (e.g., DnaK, GroES) and redox regulators were also enriched in these samples, consistent with proteotoxic and oxidative stress induced by sustained keratin metabolism (**Fig. 3d–3f and DataSet S2**). Interestingly, chemotaxis and flagellar proteins such as CheR, CheY, CheB, FlhAF and FliA were highly abundant in feather-grown cells, appearing in both cytosolic and membrane fractions. This suggests activation of chemotaxis and surface colonization mechanisms in response to recalcitrant protein substrates, a strategy likely essential for adhesion and persistence under starvation.

To integrate these protein-level responses with transcriptional data, we constructed a feather-specific protein-protein interaction (PPI) network by mapping DEGs and differentially abundant proteins to STRINGdb (**Fig. 4a and DataSet S3**). This revealed coherent modular clusters encompassing membrane proteolysis, flagellar assembly, sulfur and nitrogen metabolism, amino acid turnover, chemotaxis, ABC transporters, and redox-linked metabolic pathways (**Fig. 4a**). Such network organization underscores the coordination of substrate degradation, stress mitigation, and nutrient scavenging under environmental constraint. Regulatory hubs such as FliA (motility), FadR (fatty acid metabolism), YebC (LPS and EPS synthesis), and MarR/TetR (AcrR) repressors emerged as key nodes linking stress response and nutrient adaptation (**Fig. 4b**).

**Figure 4.**
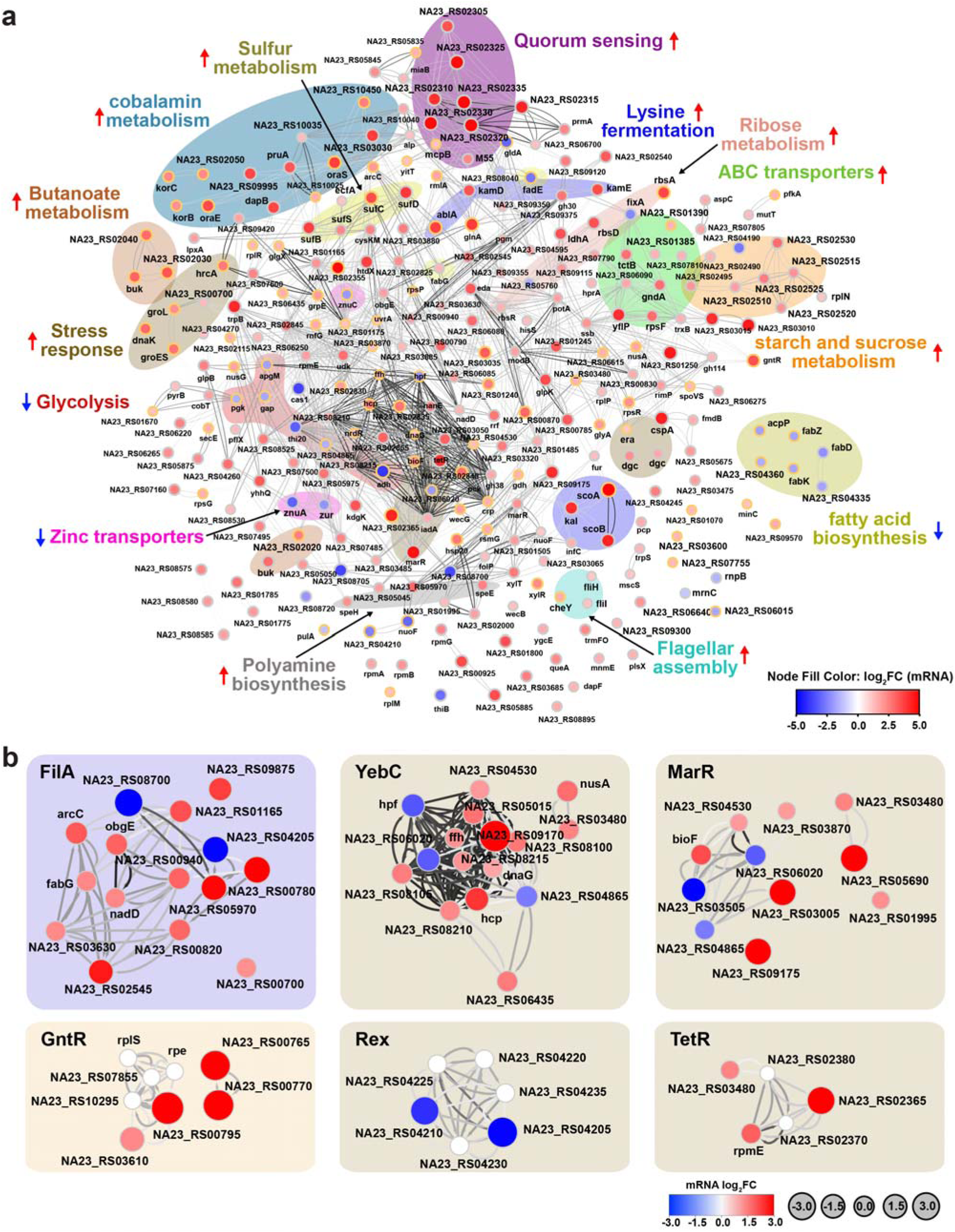
Regulatory protein-protein interaction (PPI) networks underlying starvation-induced keratinolysis in *F. islandicum* AW-1. (**a**) Global PPI network constructed from transcriptomic data comparing feather-grown versus glucose-grown cells. DEGs (log_2_FC > 1.0, *p* < 0.05, STRING combined score > 400; n = 291) were mapped to predicted interactions using STRING and visualized in Cytoscape 3.8.2 using yFiles layout plugins. Nodes fill color indicates transcript changes (red: upregulated, blue: downregulated), while black-bordered nodes denote proteins identified in the proteomic dataset. Functional clusters were annotated via KEGG pathway analysis (organism code: ‘fia’), revealing modules involved in proteolysis, motility, sulfur metabolism, and nutrient scavenging. (**b**) Subnetworks of key transcriptional regulators (FliA, YebC, MarR, GntR, Rex, and TetR) controlling stress adaptation and keratin metabolism. Edges represent known or predicted functional interactions, and node color reflects transcript abundance changes (log_2_FC).

Together, these data demonstrate that keratin degradation by *F. islandicum* AW-1 is driven by a nutrient-dependent proteomic program involving the induction of membrane-localized proteases, redox regulators, and motility machinery. The spatial colocalization of keratinases with nutrient transporters may enhance metabolic efficiency and supports survival in oligotrophic, keratin-rich environments (**DataSet. S3**).

## Keratin degradation remodels amino acid and lipid metabolism to support survival under nutrient limitation

To investigate metabolic adaptation during keratin degradation, we performed untargeted metabolomic profiling of *F. islandicum* AW-1 cultured in the presence or absence of native feathers. Time-resolved sampling from 6 to 12 hours captured key physiological transitions during growth on keratin (**Fig. 5a**). Across intracellular and extracellular fractions, 323 and 335 metabolites were detected, respectively (**DataSet S4**). Principal Component Analysis (PCA) and Partial Least Squares Discriminant Analysis (PLS-DA) showed clear separation between keratin-supplemented and control (mTF only) cultures in both intracellular and extracellular compartments, indicating substantial metabolic remodeling during keratin degradation (**Fig. 5b**).

**Figure 5.**
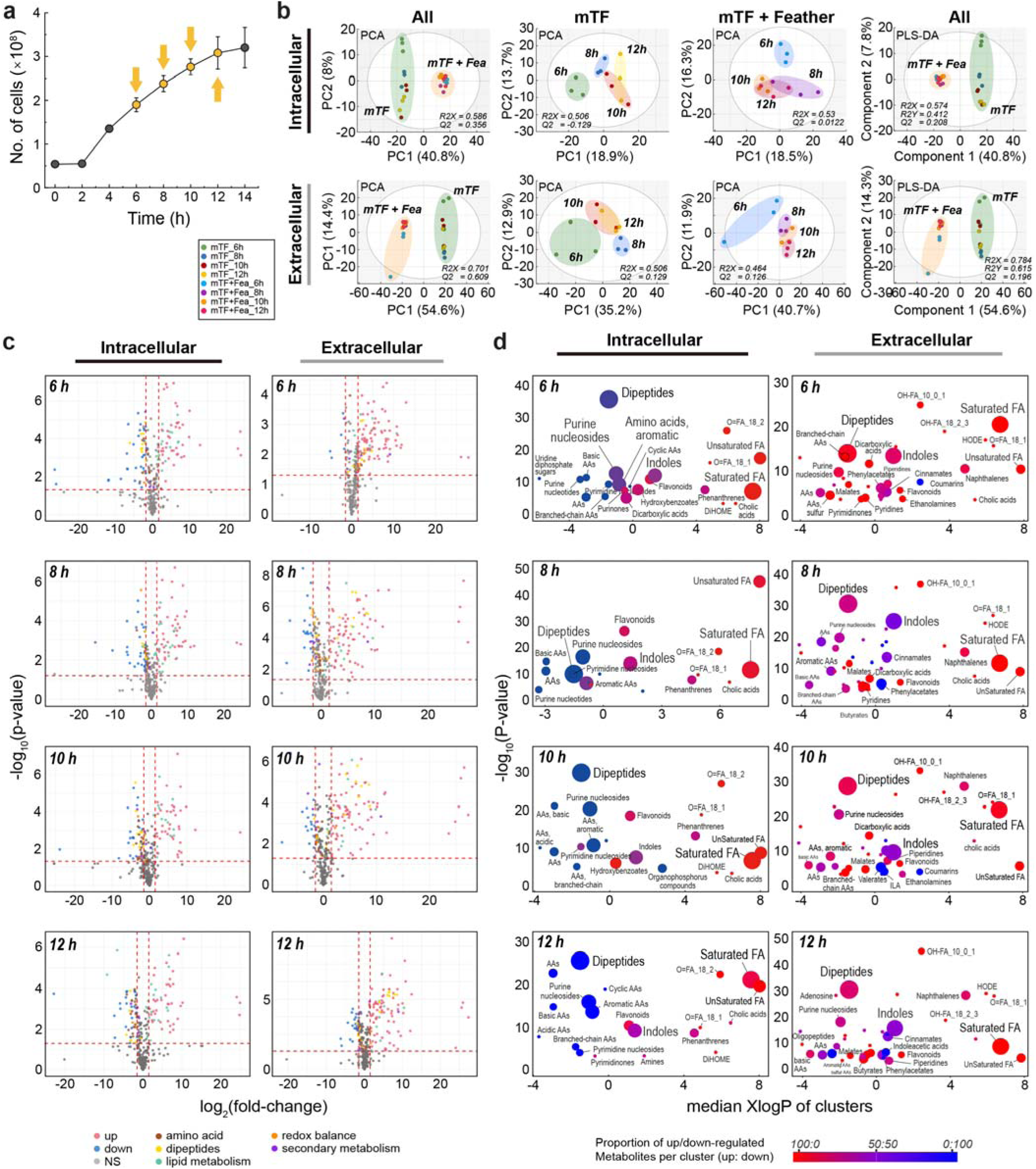
Temporal metabolomic shifts during feather degradation. (**a**). Growth curve with sampling time points for metabolomic analysis. Data represent mean ± SD (n =3). (**b**) Volcano plots of significantly altered metabolites in intracellular and extracellular fractions across time points. Metabolites with FC ≥ 2 and *p* < 0.05 were classified as upregulated (red) or downregulated (blue); grey indicates non-significant (NS) changes. **(c)** Chemical enrichment analysis of differentially abundant metabolite clusters. Bubble size indicates statistical significance (−log *p*-value), and color indicates the proportion of upregulated versus downregulated features within each cluster.

Amino acids, lipids, and metabolites related to redox balance and secondary metabolism showed significant changes in both intracellular and extracellular compartments over time (**Figs. 5c and S4**). Dipeptides (e.g., Ala-Ile, Gly-Val, Ala-Leu) and free amino acids (e.g., Val, Ser, Glu) were consistently decreased in the intracellular fraction and enriched in the extracellular fractions in feather-grown cells, consistent with proteolytic release of peptides potentially derived from keratin (**Figs. 5c–d, S4, and DataSet S4**). This pattern is consistent with transcriptomic and proteomic data, indicating elevated protease expression under feather-supplemented conditions. Increased levels of both saturated and unsaturated fatty acids (e.g., palmitoleic acid, sphingosine, nervonic acid, and oleic acid) suggest a shift in lipid metabolism, possibly supporting alternative energy production under nutrient stress.

Cluster-based chemical enrichment analysis revealed distinct temporal accumulation trends among compound classes. Dipeptides and purine nucleosides exhibited sustained depletion throughout 6–12 h, whereas saturated fatty acids progressively accumulated at later stages (10–12 h). Indoles and flavonoids showed continuous enrichment across all time points (**Figs. 5d and S4**). These metabolite profiles suggest an intial reliance on amino acid catabolism, followed by lipid mobilization and increased levels of redox-active or signaling molecules.

Structure similarity-based metabolite networks highlighted coordinated shifts in compound classes across both intracellular and extracellular environments (**Fig. 6a**). Increased levels of sulfur-containing amino acids (e.g., cysteine, methionine derivatives) and nucleotide metabolites (e.g., pyrimidine catabolites) support the integration of keratin-derived sulfur assimilation with redox regulation. Metabolic pathway mapping further revealed enrichment of ED pathway intermediates (e.g., 6PG, KDPG) and TCA cycle metabolites (e.g., succinate, α-KG, malate), consistent with transcriptome-derived evidence of metabolic rewiring for metabolic reorganization under nutrient stress (**Fig. 6b**).

**Figure 6.**
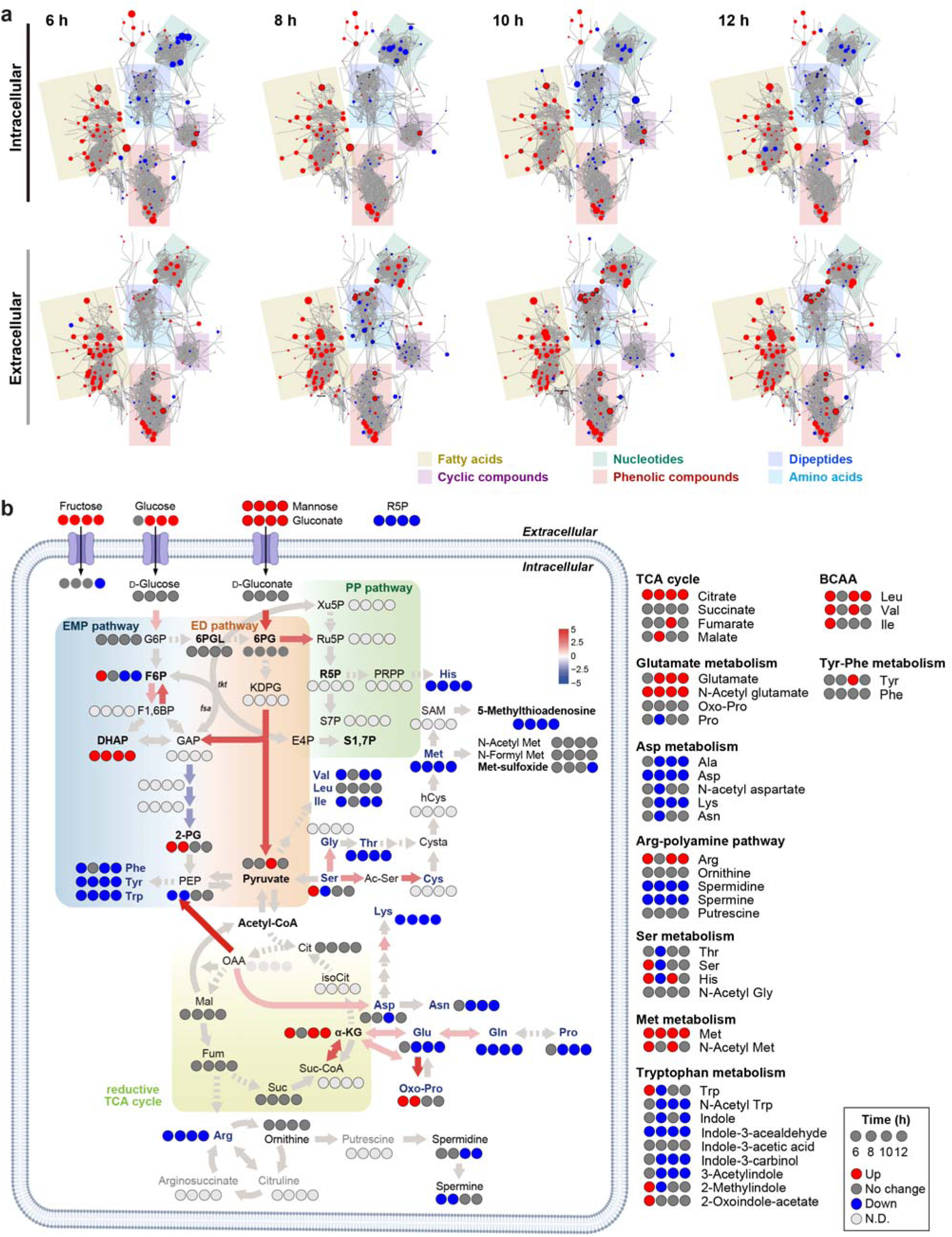
Pathway-level mapping of dynamic metabolite profiles. (**a**) Metabolite networks of intracellular (top) and extracellular (bottom) fractions clustered by chemical class with colored by fold-change direction (red: upregulated; blue: downregulated; gray: unchanged). (**b**) Temporal Log fold-change map of metabolites across central carbon and sulfur amino acid metabolism. Colored circles represent metabolites (red: increased, blue: decreased, gray: unchanged; black: undetected). Pathways include EMP, ED, and PP pathways, reductive TCA cycle, and sulfur amino acid biosynthesis. Metabolic rewiring toward the ED pathway, sulfur-containing amino acids (e.g., cysteine), and dipeptides indicates a stress-adaptive shift toward energy conservation and redox balancing.

Notably, several secondary metabolites, including indole-3-acetaldehyde and pseudouridine, exhibited dynamic changes over time (**Fig. 6 and DataSet S4**). These metabolites are associated with microbial signaling, stress adaptation, and persistence, suggesting potential activation of quorum sensing or persister cell formation under prolonged nutrient limitation.

Collectively, these metabolomic profiles support a model in which *F. islandicum* AW-1 executes a starvation-adaptive metabolic program during keratin degradation. Amino acid and lipid catabolism provide essential nutrients, while redox-active sulfur compounds and ED pathway intermediates may contribute to energy conservation and oxidative stress mitigation. In addition to nutrient acquisition, this metabolic program likely represents an ecological strategy for persistence in oligotrophic, keratin-rich environments.

## The stringent response governs chemotaxis, biofilm formation, and persister-like adaptation during keratin degradation

To elucidate the behavioral adaptation of *F. islandicum* AW-1 under nutrient-limited conditions, we examined time-resolved regulatory responses associated with starvation-induced keratin degradation. Quantitative real-time (qRT)-PCR profiling revealed early upregulation of genes related to general stress adaptive mechanisms, such as cyclic di-GMP signaling and synthesis (*dgc*, GGDEF, EAL), stringent response (*rel*, *rpoE*), and regulation of quorum sensing and biofilm formation (*luxR*). ^1, 22, 23^ (**Fig. 7a□b**).

**Figure 7.**
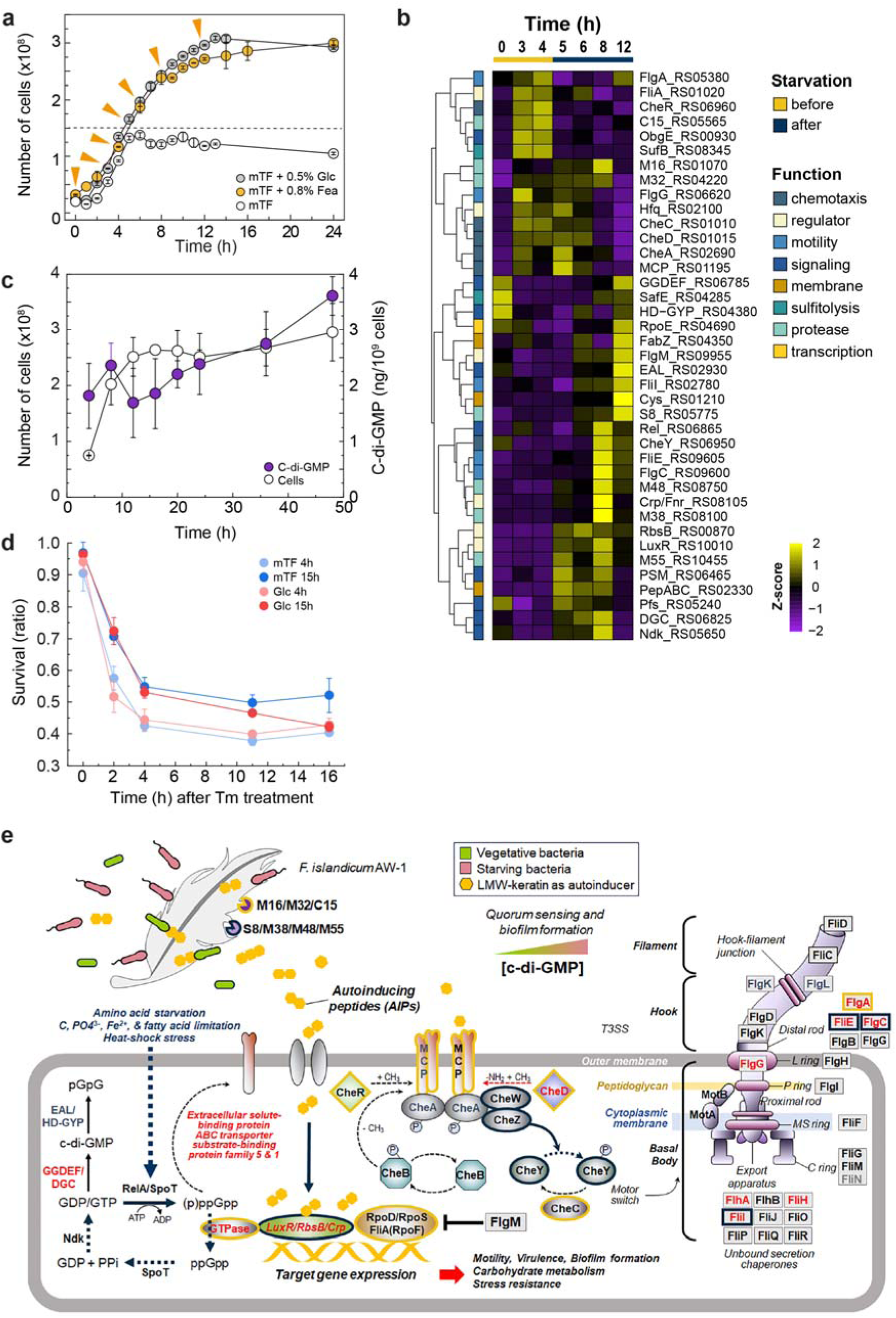
Time-resolved analysis of stringent response signaling, c-di-GMP dynamics, and antibiotic tolerance during keratin degradation by *F. islandicum* AW-1. (**a**) Growth curves in mTF medium supplemented with Glc, Fea, or no supplement. Arrows denote time points used for qRT-PCR sampling. The dashed line indicates the onset of the stationary phase in mTF medium only. (**b**) Heatmap of time-resolved gene expression profiles (0–12 h) for selected genes categorized into functional groups: chemotaxis (blue), motility (green), sulfitolysis (yellow), proteolysis (purple), transcriptional regulation (orange), and membrane-associated factors (gray). Z-scores indicate relative transcript abundance across time. (**c**) Quantification of intracellular c-di-GMP levels and viable cell counts over 48 h in feather-supplemented medium. Data are shown as mean ± SD (n = 3). (**d**) Survival assay following 200 ug/mL thiamphenicol treatment in cells representing non-starved (mTF ± Glc, exponential phase), starved (mTF + Glc, stationary phase), and deeply starved (mTF only, early death phase) physiological states. Viability was monitored over 16 h to assess antibiotic tolerance. (**e**) Schematic model of regulatory networks activated during keratin degradation under nutrient limitation. Starvation-induced signals activate the stringent response via RelA/SpoT and promote c-di-GMP synthesis via GGDEF-domain diguanylate cyclases (DGCs). Small peptides and keratin hydrolysates modulate transcriptional regulators (e.g., LuxR, Crp, RbsB), controlling chemotaxis, motility, biofilm formation, and flagellar biosynthesis. Downstream effectors (e.g., FlgM, CheY) and Type III secretion systems (T3SS) components coordinate a reverse motile-to-sessile switch, enabling persistence in keratin-rich, nutrient-depleted environments.

A central feature of this adaptation was the activation of (p)ppGpp-mediated stringent response, as indicated by transient upregulation of the RelA/SpoT homologs at 8 h (**Fig. 7b**). This timing coincided with initial nutrient depletion and likely initiated broad transcriptional reprogramming, including suppression of ribosomal genes and biosynthetic pathways, consistent with proteomic and transcriptomic evidence for energy conservation and growth arrest (**DataSet S1: Sheet 9**). In parallel, transcriptional upregulation of diguanylate cyclases (DGCs) genes containing GGDEF domains, such as NA23_RS06825, correlated with increased intracellular cyclic-di-GMP (c-di-GMP) levels between 6 10 h (**Fig. 7b and 7c**). This rise aligned with the onset of keratin hydrolysis and depletion of soluble nutrients. During this window, transcript levels of chemotaxis regulators (*cheA*, *cheR*, *cheY*) and flagellar assembly genes (*fliA*, *fliC*) also peaked, suggesting activation of motility systems to facilitate substrate localization (**Fig. 7b and 7c; DataSet S1: Sheets 1 and 9**). By 12 h, c-di-GMP levels declined, chemotaxis genes were repressed, and genes linked to adhesion and surface persistence were upregulated indicating a behavioral transition from motile to sessile states. Confocal microscopy confirmed the formation of biofilm-like microcolonies on feather surfaces (**Figs. 7c and 1h**).

Functional evidence of a persister-like state was obtained by examining antibiotic tolerance under starvation. Cells harvested from glucose-grown cultures at the exponential phase (4 h) displayed low survival upon thiamphenicol treatment (200 μg/mL), consistent with active growth and susceptibility. In contrast, cells collected from late-stationary (Glc 15 h) and prolonged starvation (mTF 15 h) conditions exhibited markedly higher survival rates despite equivalent exposure, indicating the emergence of antibiotic-tolerant phenotypes (n=6). The biphasic kill curve observed during thiamphenicol treatment mirrors classical persister cell behavior (**Fig. 7d**). Although colony formation on solid media was not feasible due to the growth limitations of *F. islandicum* AW-1, the survival patterns in liquid culture support the presence of a starvation-induced persister-like subpopulation.

Metabolite feedback has likely contributed to this regulatory shift. As keratin degradation proceeded, the extracellular accumulation of dipeptides and free amino acids may have served as signals of nutrient recovery, suppressing *spoT* expression in later phases (>12 h) and attenuating stringent signaling (**Fig. 7b**). Membrane-associated proteases, particularly S8 and M32 family enzymes, likely mediate this coupling by releasing peptide signals that integrate extracellular degradation with intracellular nutrient sensing and regulatory feedback (**Fig. 7e**).

Taken together, these results show that *F. islandicum* AW-1 orchestrates a starvation-adaptive regulatory network involving the stringent response, c-di-GMP signaling, and motility-switching. This enables a reversible transition between motile and sessile states, optimizing access to insoluble substrates and enhancing persistence under oligotrophic conditions. Notably, biofilm formation occurred in the absence of detectable exopolysaccharide production, suggesting that surface adhesion and community assembly are mediated by proteinaceous or redox-dependent interactions rather than classical EPS matrices. These findings highlight a multifaceted regulatory program that enables extremophiles like *F. islandicum* AW-1 to colonize and persist in protein-rich but nutrient-depleted environments.

## DISCUSSION

Extremophiles, such as *F. islandicum* AW-1, offer valuable insights into microbial survival strategies in oligotrophic, stress-prone environments ^24^. This study reveals that keratin degradation by *F. islandicum* AW-1 is not a simple nutrient acquisition, but rather a tightly coordinated survival strategy involving stress adaptation, metabolic rewiring, and biofilm formation (**Figs. 1 and 2**). Through multi-omics analyses, we demonstrate that keratinolysis is integrated with regulatory pathways controlling persister-like behavior (**Figs. 3 and 4**), reminiscent of survival strategies seen in modern pathogens such as *Pseudomonas* and *Mycobacterium* species ^25, 26^.

Keratin degradation is driven by membrane-associated enzymatic machinery (e.g., S8 and M48 proteases) and is further accelerated by sulfitolysis enzymes (e.g., thioredoxin, Fe-S cluster assembly proteins) under reducing conditions ^12, 18^. These enzymatic systems are embedded in a broader regulatory framework involving stringent response regulators (e.g., TetR, MarR), motility factors (e.g., CheR, FliA), and membrane remodeling genes (**Fig. 4**). This coordinated response enables *F. islandicum* AW-1 to initiate surface colonization, scavenge keratin-derived nutrients, and mitigate proteotoxic stress.

Metabolomic profiling showed significant enrichment of dipeptides, amino acids, fatty acids, and sulfur metabolites during keratin degradation. (**Figs. 5 and 6**). These compounds may serve dual roles as energy sources and signaling molecules, indicating that nutrient recycling from keratin breakdown supports both metabolism and regulatory adaptation. The enrichment of cyclic and nucleotide-derived metabolites may be associated with stringent-like stress responses although further validation is required.

Keratin degradation induces a pronounced metabolic transition from glycolysis to alternative low-energy pathways. The ED pathway is strongly upregulated, implying NADPH generation with reduced ATP expenditure, a known trait of bacteria under carbon limitation (**Fig. 2**). Upregulation of ATP citrate lyase and 2-oxoglutarate: ferredoxin oxidoreductase supports reductive TCA cycle activation, enabling CO_2_ fixation and biosynthetic precursor regeneration under oligotrophic conditions.

Sulfur metabolism is a key adaptive strategy. The Suf system facilitates both disulfide bond cleavage in keratin (sulfitolysis) and Fe-S cluster biogenesis, linking redox stress mitigation to metabolic function ^17, 18^. These adaptations parallel those seen in photosynthetic and chemolithotrophic microbes where central carbon and sulfur pathways are finely tuned under nutrient stress ^27^.

Membrane-bound proteases such as S8 and M48 remain functionally uncharacterized but are implicated in dual roles in keratin processing and cellular stress defense (**Figs. 2 and 3**). Their co-expression with cytosolic M38 and C15 peptidases, involved in protein turnover and protein homeostasis^28, 29^, supports an integrated proteostasis strategy optimized for extreme conditions ^30^.

The transition from planktonic to sessile growth on keratin surfaces suggests that *F. islandicum* AW-1 leverages biofilm-formation as a strategy for persistence under antibiotic stress ^31, 32^ and starvation ^33, 34^ (**Figs.1h and 7e**). Upregulation of flagella biosynthesis, chemotaxis, and membrane adhesion genes indicates that motility facilitates substrate localization and initial colonization (**Figs. 2, 4a, and 7b**). Despite the energetic cost, this investment likely improves substrate access and spatial niche establishment ^35^ (**Fig. 7**).

Interestingly, the observed biofilm-like state lacks canonical exopolysaccharide components, suggesting a reliance on signaling molecules—particularly low-molecular-weight (LMW) peptides derived from keratin hydrolysates—as quorum-sensing mediators (**Figs. 2 and 7**). These signals likely interact with methyl-accepting chemotaxis proteins (MCPs), modulating downstream behavior such as adhesion and persistence (**Fig. 7e**). This aligns with the elevated expression of peptide transporters and carboxypeptidases (e.g., M55, M32, M16) and supports a model in which quorum-like signaling coordinates population-level responses such as persistence and biofilm maturation ^36, 37^.

The cyclic-di-GMP signaling axis, dynamically regulated during the transition to nutrient stress, mediates this behavioral switch. Rising c-di-GMP levels coincide with sessile biofilm formation, while its transient decline aligns with increased motility and chemotaxis (**Fig. 7c**). These oscillations reflect a regulatory switch enabling environmental sensing, adhesion, and long-term persistence on keratin substrates.

These findings provide a detailed molecular portrait of starvation-induced keratin degradation in an extremophile, revealing parallels to persistence strategies in pathogenic biofilm formers ^38^. Notably, *F. islandicum* AW-1 lacks typical EPS matrices, instead using its toga-like outer membrane (a characteristic of Thermotogales) for surface interaction ^39^ (**Fig. 1d**). This adaptation, combined with flagella-driven motility, mirrors stress-induced attachment mechanisms seen in both environmental and clinical bacteria ^40^ during nutrient limitation and pathogenesis ^33, 41^.

From a biotechnological perspective, the ability of *F. islandicum* AW-1 to liquefy recalcitrant keratin under starvation offers opportunities for sustainable bioprocessing. The coordinated release of peptides, amino acids, and lipids may be exploited for value-added product recovery in waste valorization, biofertilizer production, and green chemistry application^42^.

This study provides a comprehensive model of starvation-induced keratin degradation in *F. islandicum* AW-1. By integrating multi-omics data, we uncover a survival program that couples membrane-associated proteolysis with metabolic rewiring, motility, and biofilm-associated persistence-like development. These findings offer a conceptual framework for understanding how extremophiles thrive in protein-rich but nutrient-poor environments—and may inspire new approaches to biotechnological keratin valorization and microbial survival engineering.

## MATERIALS AND METHODS

### Bacterial strains and growth conditions

*Fervidobacterium islandicum* AW-1 was cultured anaerobically in modified *Thermotoga*-*Fervidobacterium* (mTF) medium ^43^ at 70 and pH 7.0, supplemented with various carbon/nitrogen sources, as previously described ^16^. mTF medium contained: 8 g of native chicken feather, 1 g of yeast extract, 1.6 g of K_2_HPO_4_, 0.8 g of NaH_2_PO_4_·H_2_O, 0.16 g of MgSO_4_·7H_2_O, 0.1 g of NH_4_Cl, 10 mL of a vitamin solution ^44^, 10 mL of a trace element stock solution ^45^, 3 mL of 25 % (w/v) Na_2_S·9H_2_O (pH 7.0), and 1 mL of 0.1 % (w/v) resazurin per liter ^45^. Media were prepared boiled for 20 min, flushed with N_2_ gas, reduced with Na_2_S·9H_2_O (pH-adjusted), and autoclaved at 121°C for 20 min. Cultures were grown in Hungate tubes or serum bottles, sealed with black butyl rubber stoppers. Bacterial stocks were stored in 6% (v/v) dimethyl sulfoxide (DMSO) at -80 .

### Growth characterization and nutrient conditions

To assess growth under alternative nutrient conditions, mTF medium was supplemented with 5 g/L glucose, peptone, tryptone, or casein, in place of 8 g/L feathers. Seed cultures (50 mL) were pre-grown in glucose-supplemented mTF medium for 10 h at 70°C. Growth was monitored by direct cell counting using a Neubauer chamber (Marienfeld, Germany) and phase-contrast microscopy (DM750, Leica, Germany). Optical density was also measured at 600 nm (Ultrospec 8000, GE Healthcare, UK).

### Feather degradation and organic acid analysis

Feather degradation was evaluated by filtering culture broth through No. 20 filter paper, followed by drying residual feathers at 70°C overnight. Organic acid profiles were determined by HPLC (Waters 2695 Alliance, USA) using an Aminex HPX-87H column with 5 mM H_2_SO_4_ as mobile phase (35, 0.45 mL/min). Metabolites were detected by UV and RI detectors. Retention times were as follows: oxalic acid (9.22 min), citric acid (11.03), malic acid (13.18), pyruvic acid (14.82), succinic acid (16.20), lactic acid (17.68), alanine (18.18), formic acid (18.78), acetic acid (20.52), and ethanol (28.35).

### Electron and confocal microscopy

For transmission electron microscopy (TEM) and scanning electron microscopy (SEM), cells were harvested by centrifugation (10,000 × g, 10 min, 4°C), fixed overnight in 2.5% (w/v) glutaraldehyde in 0.1 M Na_2_HPO_4_/KH_2_PO_4_ buffer (pH 7.2) at 4°C, and post-fixed with 1% (w/v) OsO_4_. Samples were dehydrated through a graded ethanol series (50 to 100 %). For TEM, samples were embedded, sectioned (EM UC7/FC7, Leica), stained with 1% (w/v) ammonium molybdate, and imaged using a Hitachi HT 7700 at 80 kV. For SEM, freeze-dried samples were sputter-coated with platinum and imaged using an SU8220 (Hitachi, Japan) SEM at 15 kV.

For confocal laser scanning microscopy (CLSM), cells were stained with SYTO9 (8 μM) and propidium iodide (PI, 50 μM) for viability analysis and imaged at 400× magnification using an LSM 700 confocal microscope (Carl Zeiss). Feather cysteine residues were fluorescently labeled with CPM dye (7-diethylamino-3-(4’-maleimidylphenyl)-4-methyl-coumarin) at a 10:1 dye: thiol molar ratio, followed by 2 h incubation at room temperature. Excitation wavelengths were 480 nm (SYTO9), 538 nm (PI), and 375 nm (CPM).

### Subcellular fractionation

Cells were grown in an mTF medium supplemented with 0.5 g/L of various proteinaceous substrates or 0.8 g/L native feathers. Following the removal of feather debris by filtration (5-8 µm, Hyundai No. 20, South Korea), cells were pelleted by centrifugation (10,000 × g, 30 min, 4). The supernatant was designated as the extracellular fraction. Pelleted cells were disrupted by sonication on ice (5 min, 2 s on/5 s off, 30% amplitude). Crude lysates were centrifuged at 15,000 × g (30 min, 4°C), and the supernatant was ultra-centrifuged at 100,000 × g (2 h, 4°C) to separate cytosolic (supernatant) and membrane (pellet) fractions. Membrane proteins were solubilized in 50 mM Tris-HCl (pH 7.5) with β-dodecylmaltoside (1:1, w/w) on ice for 30 min and ultracentrifuged at 100,000 × g (30 min, 4°C) to obtain the solubilized membrane protein fraction.

### Quantification of free amino acids

Culture supernatants (50 µL) were filtered (0.45 µm), dried under vacuum, and derivatized with phenylisothiocyanate (PITC). Following a second drying step, samples were resuspended in 200 µL of solvent A (1.4 mM sodium acetate anhydrous, 0.1 % triethylamine, 6.0 % acetonitrile, pH 6.1), centrifuged at 13,000 × g (10 min, 4), and analyzed using a L-8900 amino acid analyzer (Hitachi, Japan).

For further quantification, the ninhydrin assay was also performed ^46^. Briefly, 30 µL culture broth was reacted with 150 µL cyanide-acetate buffer (10 mM KCN, 3.33 mL of NaOAc·glacial, 18 g sodium acetate trihydrate per 50 mL, pH 5.0∼5.2) and 150 µL of 3% ninhydrin solution (dissolved 2-Methoxyethanol). After incubation at 92°C for 15 min, 660 µL of isopropyl-water diluent (1:1, v/v) was added, and absorbance was measured at 570 nm.

### Enzyme activity assays

Protease activity was measured using bovine casein (Sigma-Aldrich) as substrate, following a modified Kunitz method ^47^. The reaction mixture (500 µL) contained 50 µg of enzyme and 0.2% (w/v) casein in 50 mM potassium phosphate buffer (pH 7.0). After incubation at 90 for 20 min, the reaction was terminated with 5% (w/v) trichloroacetic acid (TCA), followed by centrifugation (10,000 × g, 30 min, 4°C). The absorbance of the supernatant was measured at 280 nm. One unit (U) of protease activity was defined as the enzyme amount increasing absorbance at 280 nm of 0.01 per min.

Keratinolytic activity was measured using native feathers or soluble keratin (Chr 2, 0.2%) ^48^ as substrates. Reaction mixtures (5 mL) containing 0.5 mg of crude extract, 50 mM potassium phosphate buffer (pH 7.0), and 10 mM dithiothreitol (DTT) were incubated anaerobically at 75 . One unit (U) of keratinase activity was defined as the amount of enzyme required to produce 1 µmol of product per min. Protein concentrations were determined using a bicinchoninic acid (BCA) assay (Sigma, MO, USA) with bovine serum albumin as the standard ^49^.

### Two-chamber diffusion system

A custom anaerobic diffusion system was constructed using two 100 mL Wheaton^®^ serum bottles (223747, DWK Life Sciences) separated by a 0.20 µm cellulose nitrate membrane (Whatman, 90 mm). The semi-permeable membrane permitted the exchange of small molecules but prevented bacterial passage. The filter was sealed between rubber gaskets using a 64 mm ball joint clamp (SM.28633500, SciLab) and wrapped with Teflon tape to ensure airtight assembly. Pre-filled with 0.8% (w/v) feather-containing water, the assembled system was autoclaved, equilibrated anaerobically for 6 h, and then filled with 70 mL mTF medium per chamber. To prevent pressure imbalances and convection, manual sterile venting was performed every 12 h with a needle.

### Transcriptome analysis

Mid-exponential phase cells grown in mTF medium with 0.5 g/L glucose, peptone, tryptone, or 0.8 g/L feathers were harvested by centrifugation (10,000 × g, 30 min, 4°C), flash-frozen, and stored at -80°C. Total RNA was extracted using the RNeasy Mini Kit (QIAGEN, Germany) with on-column RNase-free DNase I treatment. All samples were ground in a mortar with liquid nitrogen prior to RNA extraction. RNA quality was verified using RNA electropherograms (Agilent 2100 Bioanalyzer) and RNA integrity numbers (RIN) scores ^50^.

For RNA-Seq, total RNA (10 µg) was used to construct sequencing libraries. mRNA was enriched using Dynabeads (Life Technologies, USA), and cDNA libraries were prepared using the TruSeq RNA Sample Prep Kit v2 (Illumina, USA). Paired-end 150-bp reads were generated on the NovaSeq 6000 platform and mapped to the *F. islandicum* AW-1 genome (GenBank accession no. NZ_CP014334.1) using Bowtie2 (v2.4.4) with default parameters. Gene-level read quantification was performed using featureCounts (v2.0.1) ^51^. Raw read counts were normalized to transcripts per kilobase million (TPM) ^52^, and differential gene expression was assessed using a customized exact test based on TPM values implemented in the *edgeR* package ^53^.

Raq RNA-Seq data have been deposited in the NCBI Sequence Read Archive (SRA) under accession numbers SRR33382689–SRR33382697 and SRX219354-7. Summary statistics of raw read counts are provided in **Table S3**. On average, ∼95% of predicted ORFs were transcribed under all tested conditions. TPM-normalized expression values, significantly differentially expressed genes (DEGs), and associated statistical parameters are integrated with the genome annotation table generated using the DNMB pipeline (https://github.com/JAEYOONSUNG/DNMB). The complete dataset is available as **DataSet 1**.

### Proteomics

#### Protein digestion and fractionation

Proteins were extracted and digested using the Filter-Aided Sample Preparation (FASP) method ^54^. Cells were lysed in SDT buffer (4% SDS, 100 mM Tris/HCl pH 7.6, 0.1 M DTT, 1 mM EDTA, 1× protease inhibitor cocktail), followed by denaturation at 95°C for 10 min and centrifugation at 16,000 × g for 5 min. Supernatants were processed on Micron YM-30 filters with UA buffer (8 M urea in 0.1 M Tris/HCl pH 8.5) to remove detergent. Alkylation was performed with 50 mM iodoacetamide in UA buffer, followed by buffer exchange with 50 mM ammonium bicarbonate by centrifugation (16,000 × g for 40 min). Proteins were digested on-filter using trypsin (1: 50, w/w) overnight at 37°C. Peptides were recovered by centrifugation at 14,000 × g for 30 min, acidified with 1% formic acid, and stored at -20°C until analysis after drying.

#### Data-dependent acquisition (DDA) and SWATH-MS analysis

Proteomic analysis was performed on a Triple-TOF™ 5600+ Mass Spectrometer (Sciex) coupled to an Eksigent NanoFlex cHiPLC system as described previously ^55^. Briefly, solvent A was composed of 0.1% formic acid in water (v/v), peptides were loaded onto a C18 trap column (0.5 mm × 200 μm) at a flow rate of 1 µL/min and separated on an analytical C18 column with a 90-min linear gradient (2–35% acetonitrile, 0.1% formic acid) at 400 nL/min. For data-dependent acquisition (DDA), survey scans (250 ms) were followed by MS/MS scans (150 ms) of the top 20 ions (precursor intensity >135, charge >1, exclusion 15 s). SWATH-MS was performed with variable 20 Da/mass windows (1 Da overlap) over the 400–1,000 m/z range. Collision energy was dynamically set per window, assuming doubly charged ions (spread ±15 eV). An accumulation time of 100 ms was used for each scan (total duty cycle of 3.1 s in high-sensitivity mode).

#### Protein identification and quantification

Raw MS data were searched using ProteinPilot (V5.0, SCIEX) against the 8460.FIAW1.2.CDS-Protein_MaxQuant_Contaminants database (2,404 entries), with trypsin specificity. ProteinPilot Software was searched with a fragment ion mass tolerance of 0.100 Da and a parent ion tolerance of 0.050 Da, applying fixed modifications for cysteine (+ 57 Da for carbamidomethylation) and biological modifications/artifacts such as methionine oxidation (+ 16 Da). Peptide and protein FDRs were controlled at <1%. SWATH-MS data (wiff files) were converted to mz5 format using ProteoWizard software and analyzed with Skyline. Peptides were quantified using three isotopic peaks per precursor and fragment, with 20 ppm mass accuracy and a 10 min retention window. Peak identity was verified by co-evolution profiles and spectral similarity. Processed data were analyzed using Perseus™ software for visualization, principal component analysis (PCA), and other statistical analyses. Proteomics data are available via ProteomeXchange (PXD027063; reviewer login; password: itBeqg1N).

### Protein interaction network analysis

STRING (ver. 11.5) interaction data for organism #2423 (*F. islandicum*) were manually curated and updated based on domain architecture and KEGG pathway mapping via InterProScan (DataSet S3). Networks were visualized in Cytoscape (ver.3.8.2.).

### Metabolomics

#### Metabolite extraction

Cells (1.0 × 10^9^) were harvested by centrifugation at 10,000 × *g* for 5 min. The cell pellets were lyophilized and disrupted with a stainless-steel ball (5 mm internal diameter) using a Mixer Mill MM400 (Retsch GmbH & Co., Haan, Germany). Intracellular metabolites were extracted using methanol:isopropanol: water (3:3:2, v/v/v, 1500 μl), followed by sonication (5 min) and centrifugation (16,100 rcf, 5 min, 4°C). Supernatants were dried in a speed vacuum concentrator. For extracellular profiling, culture media were dried and extracted using the same solvent.

#### Liquid-chromatography-orbitrap mass spectrometric analysis

The dried extracts were reconstituted in 80% methanol (50 µL) and analyzed on a Q-Exactive Plus Orbitrap (Thermo Fisher Scientific, MA, USA) coupled to an Ultimate-3000 UPLC system with a BEH C18 column (2.1 × 100 mm, 1.7 μm, Waters). Mobile phases A and B were water and acetonitrile with 0.1% formic acid (positive mode) or 0.1% acetic acid (negative mode). A solvent gradient scheme was used, starting at 0.5% organic for 0.1 min, followed by a linear increase to 80% organic over 10 min, then increase to 99.5% until 10.1 min, holding at 99.5% organic until 12 min, decreasing back until 12.1min, and holding at 0.5% organic for 3 min, for a total of 15 min. The full-scan spectra were acquired over a mass range of 80–1200 Da. MS and dd-MS2 spectra were acquired at 70,000 and 17,500 resolutions, respectively, using stepped collision energies (30, 40, and 50 eV) and a scan range of m/z 200–2000. The maximum injection time was set to 100 ms with an automatic gain control (AGC) of 10e6 charges. Data processing was performed using MS-DIAL (version 5.5.250221). Peak alignment was performed within a mass tolerance of 0.005 and alignment retention time tolerance was allowed for 0.5 min. For compound identification, accurate mass tolerance of 0.005 Da for MS1 and 0.01 Da for MS2.

#### Data processing and statistical analysis

Statistical analyses were conducted on all continuous variables (metabolites). Multivariate statistics, including principal component analysis (PCA) and partial least squares-discriminant analysis (PLS-DA), were performed using SIMCA 15 (Umetrics AB, Umea, Sweden). A chemical enrichment analysis was performed to evaluate statistical significance at the level of chemical class based on the ChemRICH program. Volcano plots were generated using GraphPad Prism 7 (GraphPad Software Inc., CA, USA). The metabolic network map was constructed based on structural similarity (Tanimoto score) and visualized by a prefuse force-directed layout using Cytoscape version 3.7.2. Multivariate empirical Bayes statistical time-series analysis (MEBA) was performed using the time-series analysis tool, web-based easy-to-use platform, MetaboAnalyst.

### Profiling of dynamic expression genes using qRT-PCR

For dynamic gene expression profiling, cDNA was synthesized from *F. islandicum* AW-1 cultures harvested at 3, 4, 5, 6, 8, and 12 hours post-inoculation, as previously described ^12^. All cDNA samples were diluted to 10 ng/µL prior to amplification. Quantitative real-time PCR (qRT-PCR) reactions (20 µL total volume) were prepared using 2× SsoAdvanced™ Universal SYBR Green Supermix (Bio-Rad), containing SYBR Green I, with 20 ng cDNA and 250 nM of each primer. Amplification was conducted on a CFX Connect™ Real-Time System (Bio-Rad) under the following cycling conditions: 95°C for 30 s, followed by 40 cycles of 95°C for 10 s and 58°C for 30 s. Primer sequences are listed in **Table S4**.

Standard curves were generated from serial dilutions of pooled cDNA to assess amplification efficiency, following the MIQE guidelines ^56^. Expression levels were normalized to the *rpo*D gene (encoding the σ RNA polymerase subunit) as the internal reference.

### Quantification of cyclic-di-GMP

Intracellular cyclic-di-GMP (c-di-GMP) levels were measured in *F. islandicum* AW-1 cultured anaerobically in mTF medium with 0.8% feathers. Samples were collected every 4 h up to 24 h and every 12 h thereafter up to 48 h. To eliminate feather debris, cultures were filtered through Whatman® Puradisc 13 syringe filters (5.0 µm, PTFE; Cytiva, USA). Cells were washed with PBS (pH 7.0), and OD600 values were measured to estimate cell density based on a previously established standard curve (6.2E+08 × (Abs.) + 2.7E+07).

Intracellular c-di-GMP was extracted using a modified heat and ethanol precipitation method, as previously described ^57^. For extraction, 1 × 10^9^ cells were fixed with 0.19% formaldehyde on ice for 10 min, washed with PBS (pH 7.0), then boiled in distilled water (100°C, 10 min). After cooling on the ice, cell lysates were mixed with ice-cold ethanol (35:65, v/v), centrifuged at 16,000 × g for 10 min at 4°C, and supernatants were pooled in three extraction rounds. Samples were flash-frozen in liquid nitrogen and lyophilized. Intracellular c-di-GMP was quantified using a commercial ELISA kit (EIAab) according to the manufacturer’s instructions.

### Observation of antibiotic tolerance

To evaluate the antibiotic tolerance of starved versus non-starved cells, *F. islandicum* AW-1 was cultivated in mTF medium with or without 0.5% (w/v) D-glucose supplementation. Cells were harvested at time points corresponding to the exponential phase (4 h), late exponential/early stationary phase (10– 12 h), and late stationary/early death phase (15–16 h) based on the mTF culture growth curve. After harvest, cell densities were normalized based on OD_600nm_ measurements and resuspended in fresh mTF medium containing 0.5% (w/v) D-glucose and 200 μg/mL thiamphenicol (Sigma-Aldrich). Cell viability was monitored by OD_600nm_ at 0, 2, 4, 10, and 16 h post-treatment. Each conditions was tested with two biological replicates and three technical replicates.

## ACKNOWLEDGMENTS

We thank Prof. Sang-Jae Lee and Jae-Won La for their technical support during the cultivation of cells and preparation of subcellular fractionated samples for keratinolytic assay and metabolomic analysis, construction of mutant strains, and differential scanning calorimetry.

## Author contributions

J.Y.S., J.Y.K., H.S.J., J.H.B. N.E.K, H.H.S., S.H.B., Y.J.L., B.C.K., D.Y.L., and D.W.L. formulated the research plan, carried out experiments, analyzed and interpreted the data, and drafted the manuscript. J.Y.S., J.Y.K., H.S.J., D.Y.L., and D.W.L. participated in the design of the study and analyzed and interpreted the data. D.Y.L. and D.W.L. conceived, planned, and supervised the study.

