## Supplementary information. for "Keratin degradation reflects a starvation survival strategy in *Fervidobacterium islandicum* AW-1"

Running Title: Keratin degradation by *Fervidobacterium islandicum* AW-1

*Corresponding authors

**Do Yup Lee**

Department of Agricultural Biotechnology, Center for Food and Bioconvergence, Research Institute for Agricultural and Life Sciences, Seoul National University, Seoul 08826, South Korea.

**Dong-Woo Lee**

Department of Biotechnology, Yonsei University, Seoul 03722, South Korea.

^‡^These authors contributed equally to this work.

Current address:

^7^Department of Environment and Energy Engineering, Gwangju Institute of Science and Technology (GIST), Gwangju 61005, South Korea.

^8^Department of Cosmetic Science & Technology, Seowon University, Cheongju 28674, South Korea.

**Table S1.** **Free amino acid concentration in the culture broth of *F. islandicum* AW-1 with native chicken feathers at 70^o^C under anaerobic conditions**

| **Feather (%)** | **Whole feather rachis** | | **0.2** | | **0.4** | | **0.8** | | **1.6** | | **3.2** | |
| --- | --- | --- | --- | --- | --- | --- | --- | --- | --- | --- | --- | --- |
| **Amino acid** | nmol | mol% | nmol | mol % | nmol | mol % | nmol | mol % | nmol | mol % | nmol | mol % |
| Asp | 0.13 | 5.6 | 17.38 | 10.21 | 18.88 | 7.73 | ND | 0.00 | 10.56 | 3.35 | 14.77 | 5.13 |
| Thr | 0.21 | 4.1 | 13.71 | 8.06 | 20.75 | 8.49 | 25.25 | 9.97 | 35.98 | 11.40 | 22.79 | 7.91 |
| Ser | 0.49 | 14.1 | 0.81 | 0.48 | 1.76 | 0.72 | 4.39 | 1.73 | 6.92 | 2.19 | 13.38 | 4.65 |
| Glu | 0.38 | 6.9 | ND^a^ | 0.00 | ND | 0.00 | ND | 0.00 | ND | 0.00 | ND | 0.00 |
| Gly | 0.54 | 13.7 | 21.44 | 12.60 | 32.62 | 13.35 | 47.07 | 18.58 | 41.63 | 13.19 | 23.72 | 8.24 |
| Ala | 1.05 | 8.7 | 64.94 | 38.17 | 88.25 | 36.11 | 89.74 | 35.42 | 115.50 | 36.60 | 108.28 | 37.60 |
| Val | 0.36 | 7.8 | 23.02 | 13.53 | 36.12 | 14.78 | 40.41 | 15.95 | 55.75 | 17.66 | 42.98 | 14.93 |
| Cys | 0.00 | 8.6 | 1.24 | 0.73 | 2.73 | 1.12 | 2.76 | 1.09 | 2.98 | 0.95 | 4.11 | 1.43 |
| Met | 0.03 | 0.1 | 3.36 | 1.97 | 3.87 | 1.58 | 4.25 | 1.68 | 6.98 | 2.21 | 10.56 | 3.67 |
| Ile | 0.11 | 3.2 | 10.43 | 6.13 | 17.49 | 7.16 | 18.60 | 7.34 | 26.52 | 8.40 | 24.89 | 8.64 |
| Leu | 0.14 | 8.3 | ND | 0.00 | ND | 0.00 | ND | 0.00 | ND | 0.00 | ND | 0.00 |
| Tyr | 0.12 | 1.4 | 5.29 | 3.11 | 8.44 | 3.45 | 9.81 | 3.87 | ND | 0.00 | 11.51 | 4.00 |
| Phe | 0.07 | 3.1 | ND | 0.00 | ND | 0.00 | ND | 0.00 | ND | 0.00 | ND | 0.00 |
| Lys | 0.18 | 0.6 | 8.53 | 5.01 | 13.49 | 5.52 | 11.04 | 4.36 | 12.77 | 4.05 | 10.98 | 3.81 |
| His | 0.40 | 0.2 | ND | 0.00 | ND | 0.00 | ND | 0.00 | ND | 0.00 | ND | 0.00 |
| Arg | 0.24 | 3.8 | ND | 0.00 | ND | 0.00 | ND | 0.00 | ND | 0.00 | ND | 0.00 |
| Pro | 1.03 | 9.8 | ND | 0.00 | ND | 0.00 | ND | 0.00 | ND | 0.00 | ND | 0.00 |
| **Total** | 5.20 | 100.0 | 170.14 | 100.00 | 244.40 | 100.00 | 253.33 | 100.00 | 315.59 | 100.00 | 287.96 | 100.00 |

Free amino acid analyses in mTF medium with varying feather concentrations after 96 h of anaerobic culture of *F. islandicum* AW-1 at 70^o^C. The amounts of amino acids in 20 µl of culture supernatant are presented.

^a^ND, not determined.

**Table S2.** **Fatty acid compositions in the cellular membrane of *F. islandicum* AW-1 grown on mTF medium supplemented with 0.5% (w/v) glucose or 0.8% (w/v) native chicken feathers**

| **Fatty acids** | **Glucose** | **Feather** |
| --- | --- | --- |
| C_10:0_ FAME^a^ | 0.97 ± 0.32 | tr^b^ |
| C_12:0_ FAME | -^c^ | tr |
| C_14:0_ FAME | 15.64 ± 1.03^d^ | 2.37 ± 0.06 |
| iso C_15:0_ FAME | - | tr |
| anteiso C_15:0_ FAME | - | tr |
| C_15:0_ FAME | - | tr |
| C_16:1_ *cis-*7 FAME | - | tr |
| C_16:1_ *cis*-9 FAME | - | 2.07 ± 0.08 |
| C_16:0_ FAME | 79.06 ± 0.09 | 44.15 ± 0.21 |
| C_17:0_ FAME | - | 1.84 ± 0.05 |
| C_18:2_ *cis*-9, 12 FAME | - | 1.54 ± 0.07 |
| C_18:1_ *cis*-9 FAME | 1.13 ± 0.19 | 23.49 ± 0.08 |
| C_18:0_ FAME | 3.28 ± 0.93 | 13.59 ± 0.47 |
| C_19:0_ FAME | - | tr |
| C_20:1_ *cis*-11 FAME | - | tr |
| C_20:0_ FAME | - | 2.25 ± 0.32 |
| Summed feature 2^e^ | - | tr |
| Summed feature 10^f^ | - | 2.52 ± 0.08 |
| Summed feature 11^g^ | - | tr |
| Summed feature 12^h^ | - | 1.11 ± 0.01 |

^a^FAME, fatty acid methyl ester

^b^tr, trace (<1%)

^c^-, not detected.

^d^Data are expressed as means ± standard deviations (SDs).

^e^Summed feature 2 comprises C_12:0_ 3-0H FAME and/or C_13:0_ dimethyl acetal (DMA).

^f^Summed feature 10 comprises C_18:1_ c11/t9/t6 FAME and/or UN 17.834.

^g^Summed feature 11 comprises iso-C_17:0_ 30H FAME and/or C_18:2_ DMA.

^h^Summed feature 12 comprises UN 18.622 and/or iso-C_19:0_ FAME.

**Table S3. A summary of raw reads of RNA-Seq data**

| **Sample ID** | **Total read bases (bp)** | **Total reads** | **Sequencing platform** | **GC (%)** | **AT (%)** | **Q20 (%)** | **Q30 (%)** |
| --- | --- | --- | --- | --- | --- | --- | --- |
| FAW1-Fea-1 | 967,574,302 | 18,897,624 | Illumina HiSeq 2500 | 43.0 | 57.0 | 96.3 | 92.7 |
| FAW1-Fea-2 | 2,583,525,864 | 25,579,464 | Illumina NovaSeq 6000 | 44.7 | 55.3 | 98.6 | 95.5 |
| FAW1-Fea-3 | 2,495,933,210 | 24,712,210 | Illumina NovaSeq 6000 | 45.9 | 54.1 | 98.7 | 95.5 |
| FAW1-Pep-1 | 1,665,824,893 | 32,705,112 | Illumina HiSeq 2500 | 43.2 | 56.8 | 96.5 | 93.0 |
| FAW1-Pep-2 | 3,687,524,746 | 36,510,146 | Illumina NovaSeq 6000 | 44.4 | 55.6 | 98.9 | 96.1 |
| FAW1-Pep-3 | 2,324,358,450 | 23,013,450 | Illumina NovaSeq 6000 | 44.1 | 55.9 | 98.6 | 95.3 |
| FAW1-Trp-1 | 1,100,011,425 | 21,597,348 | Illumina HiSeq 2500 | 43.8 | 56.2 | 96.4 | 92.9 |
| FAW1-Trp-2 | 2,226,273,916 | 22,042,316 | Illumina NovaSeq 6000 | 45.1 | 54.9 | 98.6 | 95.3 |
| FAW1-Trp-3 | 2,041,061,732 | 20,208,532 | Illumina NovaSeq 6000 | 44.9 | 55.1 | 98.7 | 95.5 |
| FAW1-Glc-1 | 1,161,356,801 | 22,801,919 | Illumina HiSeq 2500 | 43.1 | 56.9 | 96.4 | 92.9 |
| FAW1-Glc-2 | 2,322,133,218 | 22,991,418 | Illumina NovaSeq 6000 | 46.5 | 53.5 | 98.3 | 94.6 |
| FAW1-Glc-3 | 3,665,953,166 | 36,296,566 | Illumina NovaSeq 6000 | 46.7 | 53.3 | 98.5 | 95.0 |
| FAW1-Fea_12h-1 | 2,260,657,750 | 22,382,750 | Illumina NovaSeq 6000 | 44.6 | 55.4 | 97.7 | 93.8 |
| FAW1-Fea_12h-2 | 3,143,896,892 | 31,127,692 | Illumina NovaSeq 6000 | 50.1 | 49.9 | 97.6 | 93.8 |
| FAW1-Fea_12h-3 | 3,161,199,808 | 31,299,008 | Illumina NovaSeq 6000 | 45.6 | 54.4 | 97.7 | 93.8 |

**Table S4. Primers used for qRT-PCR analysis**

| **Gene name** | **Sequence (5'-3')** | **Tm (℃)** |
| --- | --- | --- |
| RS02690 (*che*A) | AGAGGGTGAACCACAAAAGG | 59.0 |
|  | CTTCGTGCAATCACAAGCTC | 59.6 |
| RS01010 (*che*C) | ATCTGTCCCGCAAGTAAAGG | 59.2 |
|  | TTTGGGTCGAAGATCAGCAG | 61.3 |
| RS01015 (*che*D) | CACGGGTGTTAATCTTGTGG | 58.9 |
|  | ACCGCTTCTACATTCCTTGC | 59.3 |
| RS06960 (*che*R) | ACGGCACAATCTTCTCCAAG | 60.3 |
|  | CGAATAATAGTCCGCCAACG | 60.5 |
| RS06950 (*che*Y) | CGAAAGATGGGATGGAACAG | 60.5 |
|  | ACTCCACAGCCTTTTCAACG | 60.3 |
| RS01195 (*mcp*) | AAACTCGCAACTGCCTTGAG | 60.6 |
|  | TCTTCGATTGTCGCACTGAC | 60.0 |
| RS01020 (*fli*A) | TGCCAAAAACCAGCGACTAC | 61.2 |
|  | CTTCTTCGTCCGAACCAAAC | 59.7 |
| RS09605 (*fli*E) | GGTGGGGTTAATCCGTTAAG | 58.3 |
|  | CGGTCAACTTTTCCACGTTC | 60.5 |
| RS02780 (*fli*I) | ATCGGTGAACCTCCAACAAC | 59.8 |
|  | ATCGGCTTCGACAAGAACTG | 60.4 |
| RS05380 (*flg*A) | TCGTTGCATACGTTCCTGTC | 59.7 |
|  | GAGCCCTTTGTTGTGCTTTC | 59.9 |
| RS09600 (*flg*C) | GGCGCAAAGGTTCAGAATAG | 59.8 |
|  | ACCAGAGTTTTCACGCTTGC | 60.4 |
| RS06620 (*flg*G) | TTTGGCTATCAGTGGTGACG | 59.7 |
|  | AGAGATACCGCGTTTTGTGG | 60.1 |
| RS09955 (*flg*M) | CGGAGGTCAAAGGAAAAACC | 60.8 |
|  | ATACTCGGCCACTTTCCTTC | 58.3 |
| RS06825 (*dgc*) | TCTTTCTTGCCCTCAGGTTC | 59.4 |
|  | CTGCTGTATGCACCGGTTAG | 59.4 |
| RS00930 (*obg*E) | CGCAGAGAATGGAGAAAACG | 60.9 |
|  | TATTTCCCAGGCTCGTCAAG | 60.2 |
| RS06865 (*rel*) | TCCGAGTTGGAGGATTTGAG | 60.2 |
|  | TGCTCTTGGAACGATGTCTG | 60.0 |
| RS05650 (*ndk*) | AGGGAAGCCGTTTTACCAAG | 60.5 |
|  | TGCTCCGACTATGTGCCTTAC | 60.3 |
| RS06785 (*dgc*) | AGATACGGAGGAGACGAGTTTG | 59.8 |
|  | TCGACTAAGGTTTGCCCTTG | 60.2 |
| RS02930 (*eal*) | TCACGAGCAAAGTCGTCAAG | 60.2 |
|  | TCTTCAGCTCGCCATTGTAG | 59.2 |
| RS04380 (*hd-gyp*) | TCACCTTTCTCATCGGCTTC | 60.3 |
|  | TCAACGTATTCACCGGTCTC | 58.6 |
| RS05240 (*pfs*) | GCGTTCGCAGTTGATATGG | 60.2 |
|  | TAATGGCAGAGGGCAAACTC | 60.2 |
| RS06465 (*psm*) | AATCCGGAGGACTTGAAAGC | 60.6 |
|  | CCATTGCACTTGACCATGAC | 60.0 |
| RS04350 (*fab*Z) | CGAGGCAAGGTTCAAGAAAG | 60.0 |
|  | TCACCGACCTTAGCTTTTCC | 59.3 |
| RS02100 (*hfq*) | GGGGATTGTTAGGTCGTTTG | 59.3 |
|  | GTTCCGATTCTTGCTCTTCG | 60.0 |
| RS00870 (*rbs*B) | TCTCCGTCTTTCTTCCTTGC | 60.0 |
|  | TCGCTGCAGATAGTGCAAAG | 60.3 |
| RS10010 (*lux*R) | CGAAGAGATCCTCGAAATGG | 59.8 |
|  | TCTCCGTCTTTCTTCCTTGC | 59.5 |
| RS08105 (*crp*/*fnr*R) | CAGCGGGAACGAAAATCTAC | 59.7 |
|  | GATGGCAGGTTGAAAAGAGC | 59.8 |
| RS00485 (*rpo*D) | TGTGGGTGGTGAGAAGATTG | 59.5 |
|  | AGCTTCTTCATCACCCTTCG | 59.4 |

Flagellar biosynthesis (9 genes), chemotaxis (6 genes), ppGpp and C-di-GMP signaling (8 genes), autoinducer production (3 genes), and transcriptional regulation (4 genes)-related genes were analyzed for *F. islandicum* AW-1 after 3, 4, 5, 6, 8, and 12 hours of anaerobic culture in mTF+0.8% fea medium at 70°C. The Tm values were calculated using Primer3Plus.

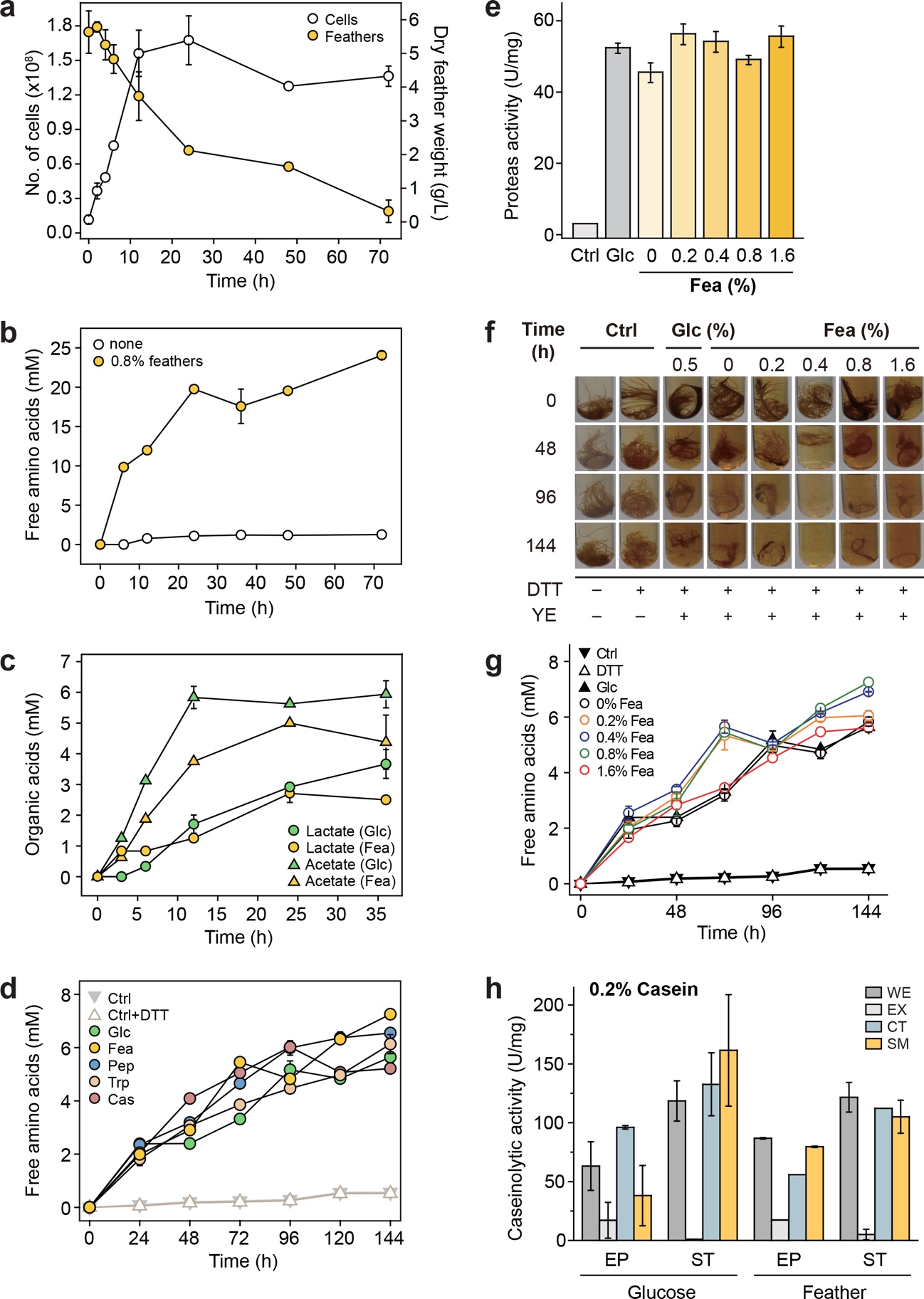

**Figure S1. Nutrient- and redox-dependent keratin degradation by *F. islandicum* AW-1. (a)** Time-course of cell growth and feather degradation in mTF medium supplemented with 0.8% native chicken feathers at 70℃ and pH 7.0. Feather biomass was quantified by dry weight. (**b**) Accumulation of free amino acids in cultures with 0.8% feathers (yellow) or no supplement (open circles). (**c**) Organic acid production during growth on 0.8% feathers (yellow) vs. 0.5% glucose (green). (**d**) Free amino acid release from cells grown on various nutrient conditions (Glc, Fea, Pep, Trp, Cas), with or without 10 mM DTT. (**e**) Total protease activity in crude extracts from cells grown on Glc or increasing concentrations of Fea (0.2–1.6%, w/v). Data represent means ± SD (n=3). (**f**) **Time-course of feather degradation using crude extracts from glucose- or feather-grown cells, with or without DTT** supplementation. Reactions were carried out at 75 °C and pH 7.0 for 144 h. (**g**) Keratinolytic activity at different feather concentrations, measured by the release of free amino acids from native feathers over 144 h at 75°C. (**h**) **Caseinolytic activity of** subcellular fractions (WE, EX, CT, SM) from exponential (EP) and stationary (ST) phase cells **grown on Glc or Fea.** Caseinolytic activity was conducted using 0.2% casein at 90℃ and pH 7.0 for 20 min.

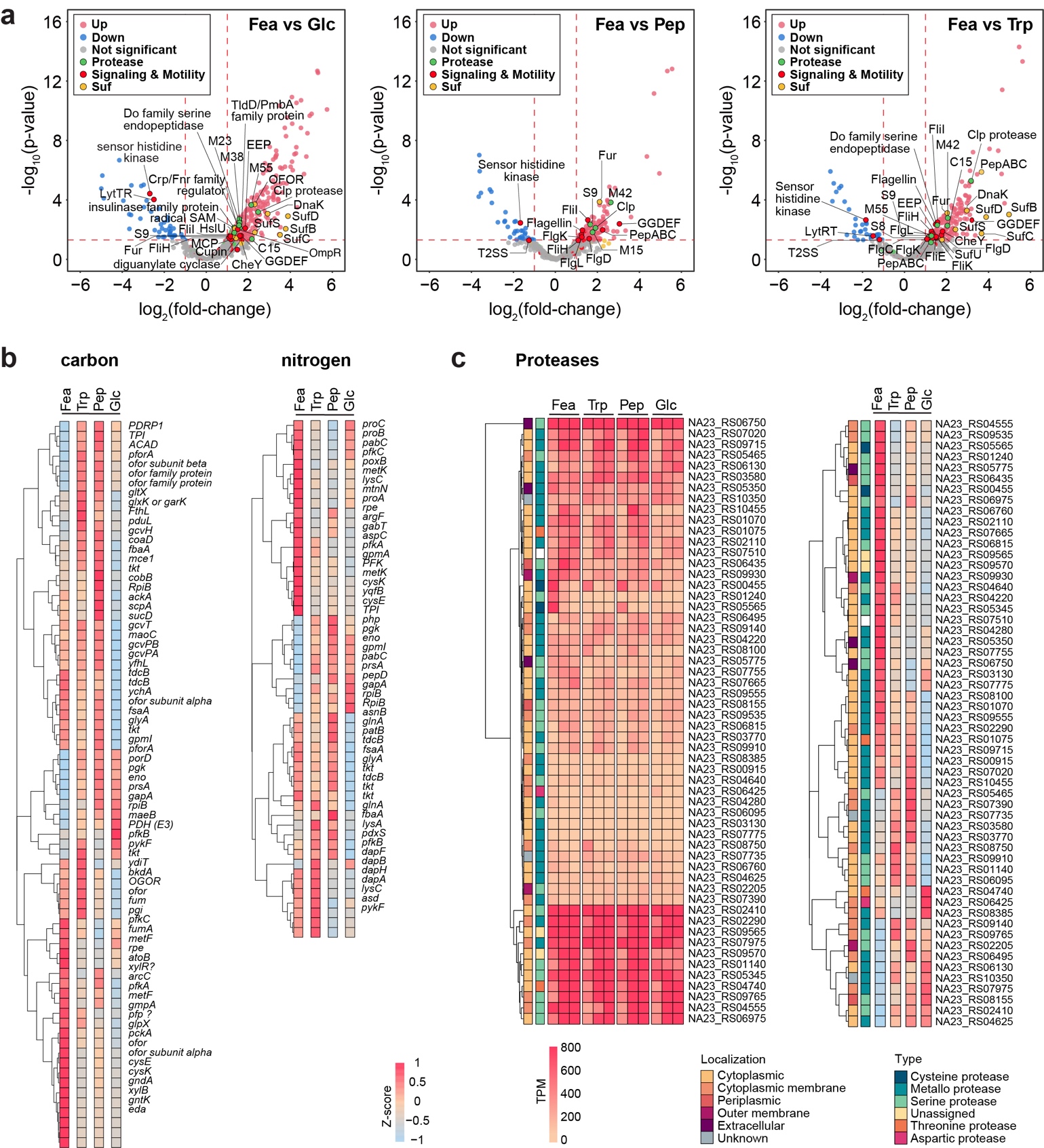

**Figure S2. Nutrient-specific transcriptomic responses and protease gene expression in *F. islandicum* AW-1.** (**a**) Volcano plots comparing DEGs from cells grown on feathers (Fea), peptone (Pep), or tryptone (Trp) relative to glucose (Glc). All samples are biologically replicated in triplicate. Red dots indicate significantly upregulated genes (≥ 2.0 fold, *p* ≤ 0.05). Blue dots indicate significantly downregulated genes (≤ -2.0 fold, *p* ≤ 0.05). Grey dots represent non-significant genes (*p >* 0.05). Functionally relevant genes related to proteases, signaling & motility, and sulfur metabolism (Suf) are highlighted. (**b**) Heatmaps of DEGs involved in nitrogen and carbon metabolism. Genes involved in nitrogen metabolism (left) and carbon metabolism (right) are clustered based on their expression levels. Red indicates high expression, while blue represents low expression (Z-score normalized). (**c**) Heatmaps of 57 protease-encoding genes under different nutrient conditions. The heatmap on the left shows the expression of protease genes categorized by cellular localization (cytoplasmic, periplasmic, outer membrane, extracellular). The heatmap on the right classifies proteases by enzyme type, including cysteine, metalloprotease, serine, and threonine proteases. Expression levels are shown in transcripts per million (TPM).

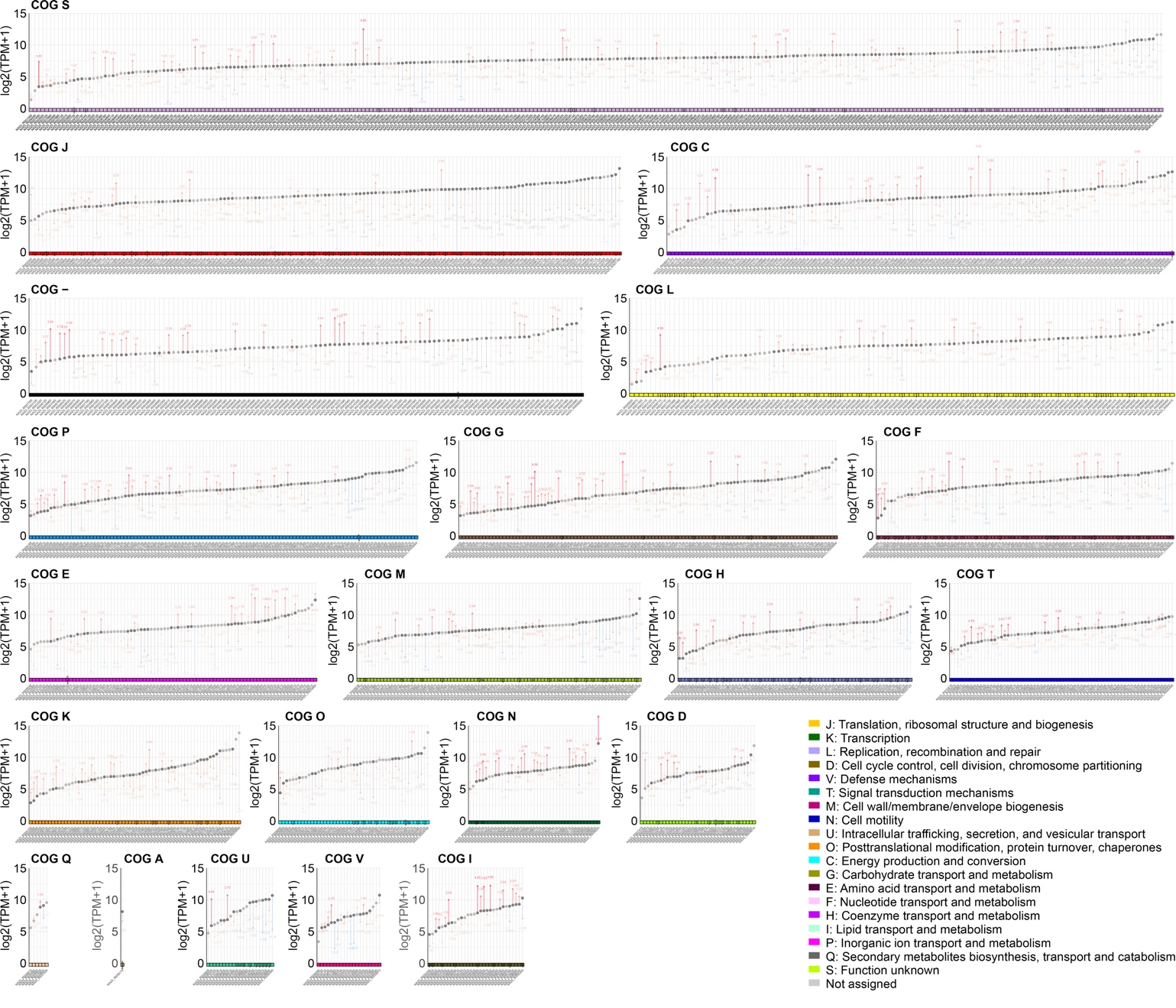

**Figure S3. Functional category-specific transcriptomic changes during late-stage keratin degradation.** COG-based expression profiles comparing Fea-grown cells at 12 h vs. 8 h. Genes are grouped by COG category, and their expression is shown as log₂(TPM + 1). Vertical arrows represent relative transcript changes at 12 h compared to 8 h: red for upregulated genes, blue for downregulated. Significant induction is observed in stress response (O), translation (J), membrane biogenesis (M), and energy metabolism (E, G, P), indicating a transition to a stress-adaptive state during prolonged feather degradation.

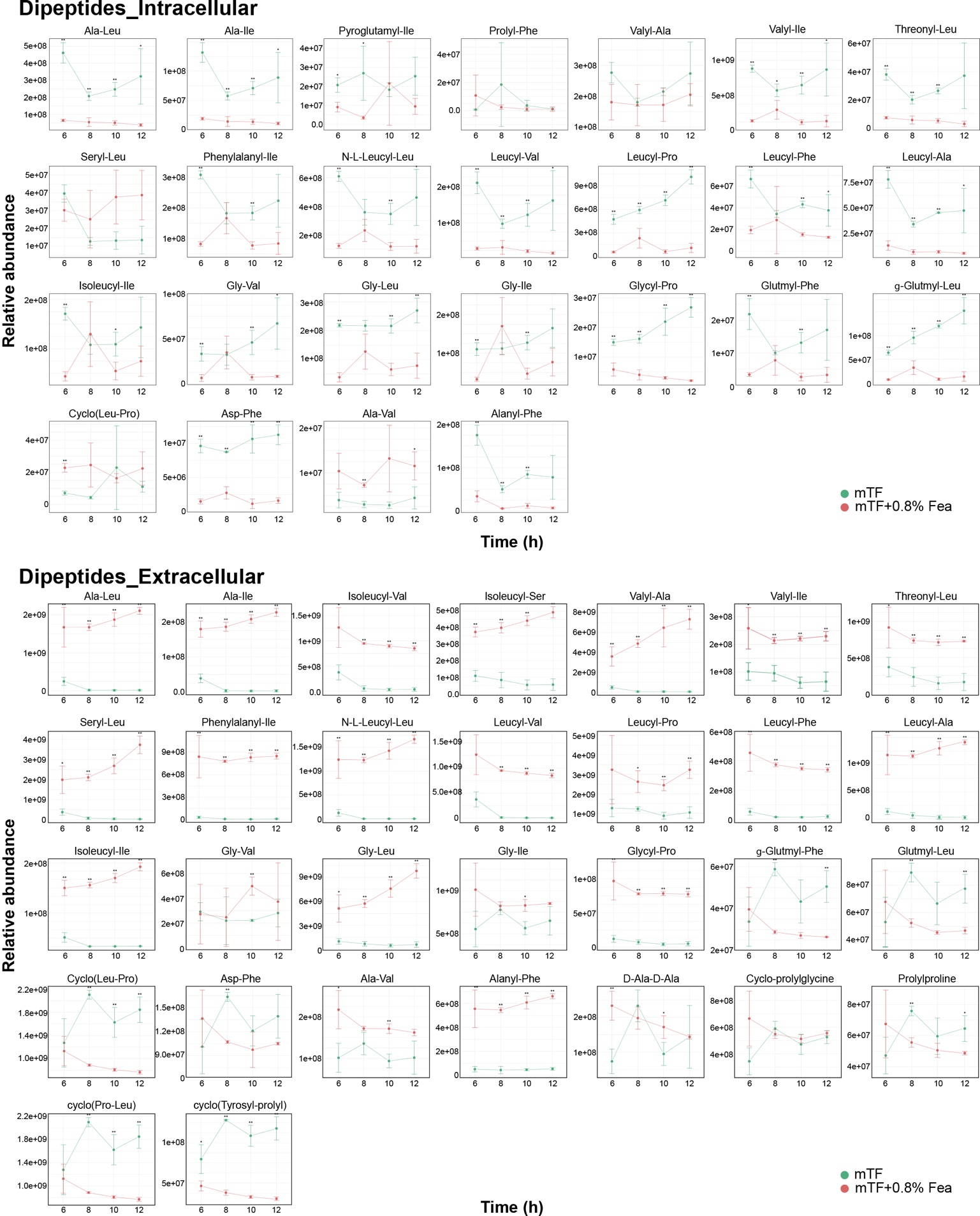

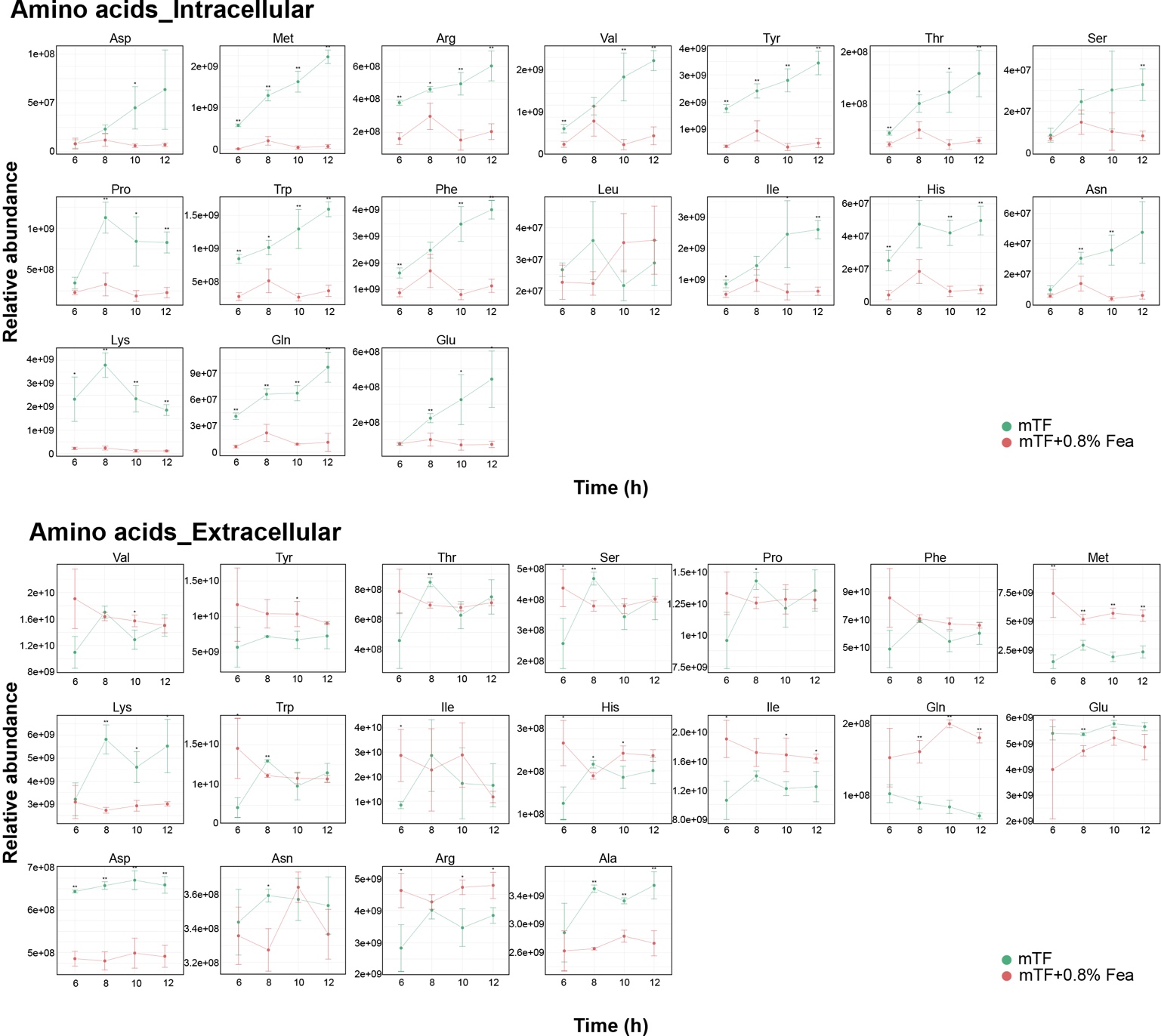

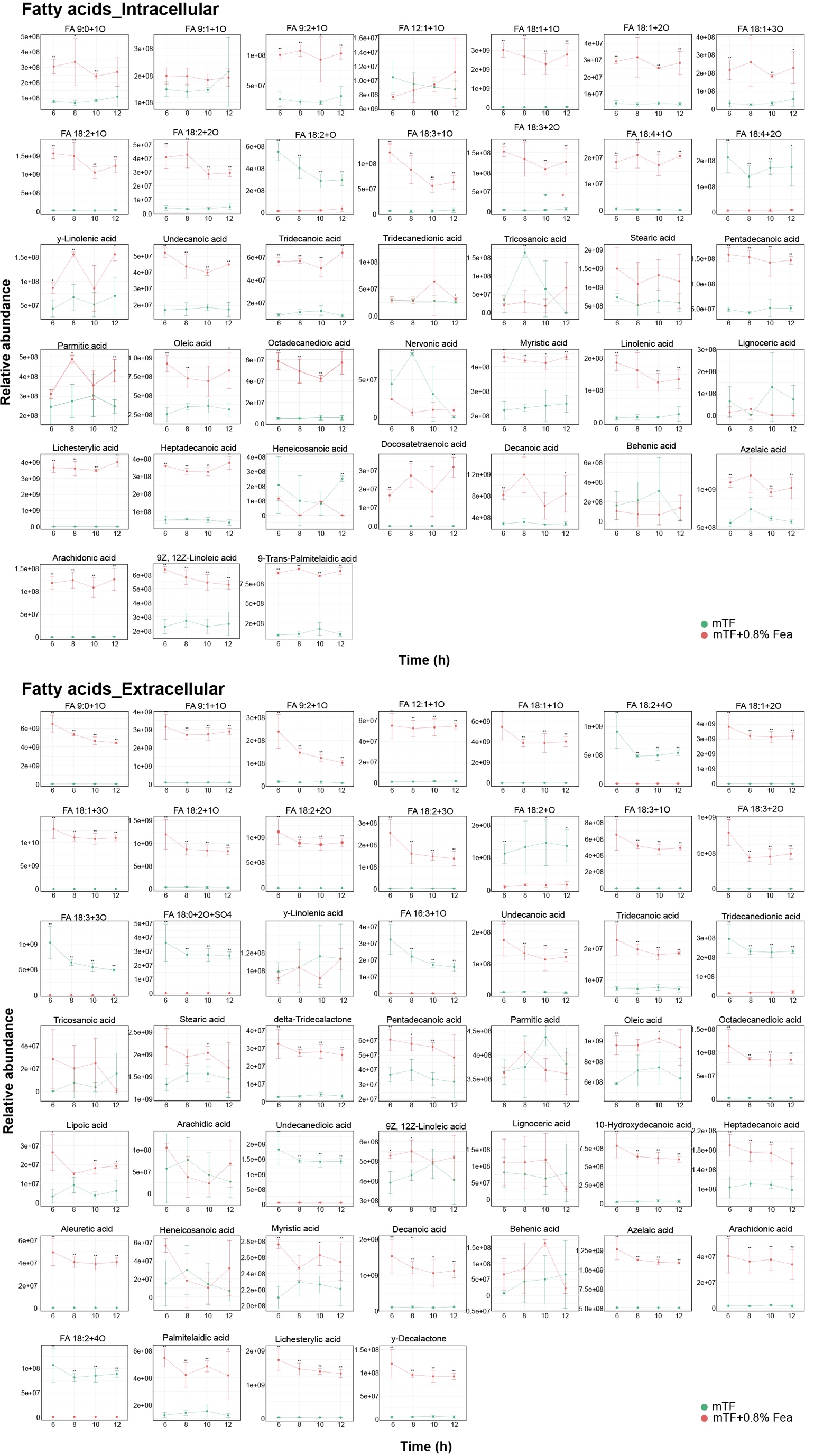

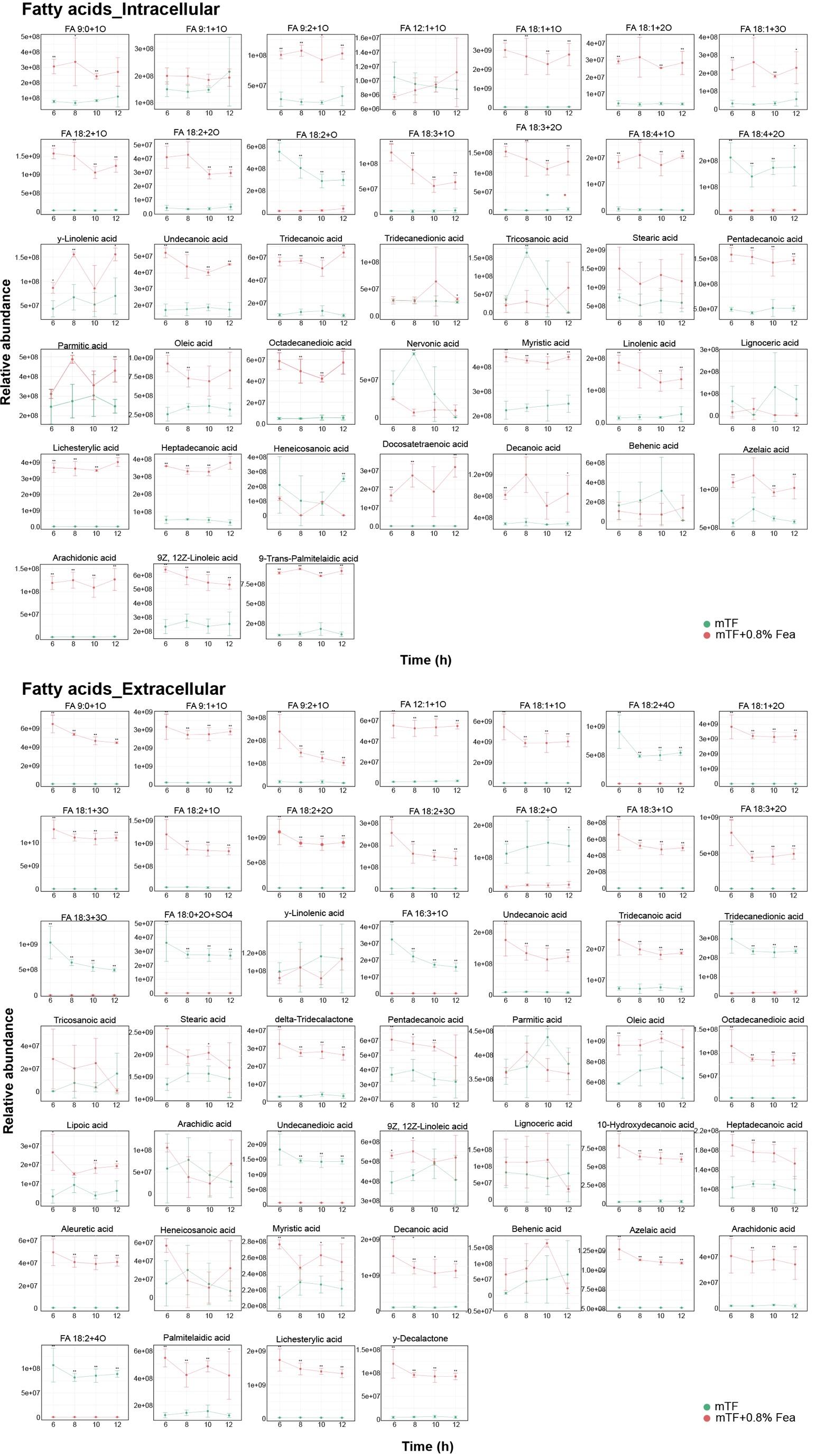

**Figure S4. Temporal metabolomic profiling of dipeptides, amino acids, and fatty acids during feather degradation by F. islandicum AW-1.** Time-resolved intracellular and extracellular abundances of dipeptides, amino acids, and fatty acids measured at 6, 8, 10, and 12 h during growth in mTF medium supplemented with or without 0.8% (w/v) feathers. Each panel displays relative abundances (log scale) of individual metabolites, grouped by compound class and compartment (intra-/extracellular). Data are shown as mean ± SD (n = 3). Metabolites were selected based on significant temporal changes or nutrient-dependent differences (adjusted *p* < 0.05).
